# Extending the IICR to multiple genomes and identification of limitations of some demographic inferential methods

**DOI:** 10.1101/2024.08.16.608273

**Authors:** Lounès Chikhi, Willy Rodríguez, Cyriel Paris, Marine Ha-Shan, Alexane Jouniaux, Armando Arredondo, Camille Noûs, Simona Grusea, Josué Corujo, Inês Lourenço, Simon Boitard, Olivier Mazet

## Abstract

Reconstructing the demographic history of populations and species is one of the greatest challenges facing population geneticists. [50] introduced, for a sample of size *k* = 2 haploid genomes, a time- and sample-dependent parameter which they called the IICR (inverse instantaneous coalescence rate). Here we extend their work to larger sample sizes and focus on *T*_*k*_, the time to the first coalescence event in a haploid sample of size *k* where *k* ≥ 2. We define the IICR_*k*_ as the Inverse Instantaneous Coalescence Rate among *k* lineages. We show that (i) under a panmictic population 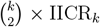 is equivalent to *N*_*e*_, (ii) the IICR_*k*_ can be obtained by either simulating *T*_*k*_ values or by using the *Q*-matrix approach of [61] and we provide the corresponding Python and R scripts. We then study the properties of the 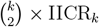 under a limited set of *n*-island and stepping-stone models. We show that (iii) in structured models the 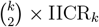 is dependent on the sample size and on the sampling scheme, even when the genomes are sampled in the same deme. For instance, we find that 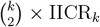 plots for individuals sampled in the same deme will be shifted towards recent times with a lower plateau as *k* increases. We thus show that (iv) the 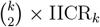 cannot be used to represent “the demographic history” in a general sense, (v) the 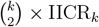 can be estimated from real or simulated genomic data using the PSMC/MSMC methods [44, 65] (vi) the MSMC2 method produces smoother curves that infer something that is not the 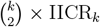, but are close to the 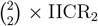 in the recent past when all samples are obtained from the same deme. Altogether we argue that the PSMC, MSMC and MSMC2 plots are not expected to be identical even when the genomes are sampled from the same deme, that none can be said to represent the “demographic history of populations” and that they should be interpreted with care. We suggest that the PSMC, MSMC and MSMC2 could be used together with the 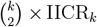 to identify the signature of population structure, and to develop new strategies for model choice.

## 1 Introduction

Population geneticists have witnessed major changes in the way they are obtaining, analysing and interpreting genetic data in the last decades [81, 85, 71]. It has become increasingly easy to obtain tens or hundreds of thousands of genetic markers and full genomes even for non model species [28, 86, 87, 23, 83, 30, 57, 74, 75, 5] including those for which only museum and archaeological samples are available [24, 51, 60]. In parallel to the increase in data, sophisticated statistical methods have been developed that allow population geneticists to process, filter and analyse these data for demographic inference [44, 22, 65, 46, 13, 41, 71, 81, 4, 14]. Major technical and technological progress is thus currently taking place including in the simulation of genetic and genomic data [38, 42, 22]. However, the interpretation of genomic data remains difficult and still requires significant amounts of testing and validation, and a better understanding of the properties of genomic data under different demographic models.

One major tradition of population genetic research has focused on the demographic history of populations, where the phrase “demographic history” is interpreted in terms of effective population size (*N*_*e*_) changes. This tradition has provided important insights into the recent evolutionary history of species including humans and other primates [73, 70, 62, 7, 26, 58, 83, 30] and is the basis of several important inference methods [7, 44, 65, 46, 15, 13]. However, it may also have led to potentially disputable conclusions as noted by several authors [25, 6, 17, 19, 20, 77] and there is an increasing recognition that population structure can generate spurious signals of population size change [79, 80, 6, 52, 72, 56, 32, 55, 49, 50, 61, 16, 29, 4]. We know now that a history of population size changes may be inferred with misleadingly great confidence simply because the population was structured. This is the case even if the population size never changed or changed in the direction opposite to that inferred. In that context, genomic data may lead to an overconfidence in parameter estimates that may have little to do with the actual history of populations [17, 19, 20, 49, 50, 16, 77]. It seems thus important to develop methods or theoretical frameworks that integrate population structure during the inferential process [16, 81, 4, 77]. Here we extend the work of [50], [16] and [61] on the IICR (inverse instantaneous coalescence rate) to *k >* 2 genomes. We study some of its properties and discuss how it could be used to improve our understanding of structured populations. For the sake of consistency, the original IICR of [50] will be identified as IICR_2_ in the rest of the manuscript. The IICR_2_ is a time- and sample-dependent parameter originally defined for a sample size of *k* = 2 (two haploids or one diploid). In a panmictic population, the IICR_2_ is the equivalent of the coalescent effective population size, *N*_*e*_ [37, 69]. In structured populations, however, the IICR_2_ can be strongly disconnected from changes in the total number of individuals or from *N*_*e*_ estimates (see [16] for a short discussion on the relation between the IICR_2_ and the concept of *N*_*e*_). The IICR_2_ is a function of the distribution of coalescence times and is thus a function of the demographic model assumed and of the geographical and temporal location of the samples [50, 16, 29, 61]. This can be loosely written as IICR_2_ = *f*_Mod_(*t, k* = 2, [*v*]), where Mod represents the demographic model, *k* = 2 is the number of haploid genomes, [*v*] is a vector representing the sampling configuration and *t* represents time in units of 2*N* generations, starting at the sampling time of the most recent sample, and *N* is the number of haploid genomes (see next section).

Currently, the IICR_2_ can be computed for any model for which the *T*_2_ can be derived analytically or numerically, or simulated [50, 16, 61]. With real data, it can be estimated by the PSMC (pairwise sequentially Markovian coalescent) method of [44]. For larger *k* values, no theoretical work as been developed even if the MSMC method of [65] should be able to estimate it for genomic data. As we will show here, a simulation-based approach can be used to approximate the IICR_*k*_ for any scenario for which *T*_*k*_ values can be obtained as in [16] for *T*_2_ values. We stress that the IICR_*k*_ needs to be rescaled because the coalescence rate varies with both *N* and *k* in a panmictic population such that the coalescence rate is larger for increasing *k* values by a factor of 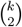. This means that it is 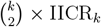 that is the equivalent of *N*_*e*_ in a panmictic population. Finally, we note that we can also obtain an exact computation of the IICR_*k*_ by extending the transition matrix approach of [61] that was developed for *k* = 2.

In practice, we thus (i) extend the IICR_2_ to the case where the number of haploid genomes is *k* ≥ 2 and define the IICR_*k*_ as the inverse of the first coalescence rate for a sample of size *k*; (ii) apply this approach to structured models of population genetics for which we predict and compare the 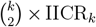 obtained for different *k* values and sampling schemes, (iii) compare these results to those obtained using the MSMC method of [65] for genomic data generated under the same models, (iv) show that the PSMC and MSMC methods are in agreement with our theoretical results, (v) show that the two methods are subject to significant stochasticity with plots that can differ significantly from the expected 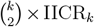 plot, (vi) show that the MSMC2 method exhibits less stochasticity than the PSMC and MSMC methods but does not generally estimates the 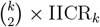 (vii) discuss how these 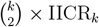 plots could be used for model choice or model exclusion, (viii) provide Python and R scripts to compute and plot the 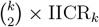, (ix) discuss the limitations of the phrase “demographic history”.

## 2 Methods

### 2.1 Definition of the IICR_*k*_

Let us assume that we have a sample of *k* genes at the present (time *t* = 0). We call *T*_*k*_ the time to the first coalescence event between any pair of genes in the sample, where time is scaled by *N*_0_, the number of haploid genomes at *t* = 0. *T*_*k*_ is a random variable that takes values in R_+_ and whose pdf (probability density function) is a function of the evolutionary history of the population or species of interest. [27] showed that in a panmictic population where the size at a given time (denoted *N* (*t*)) varies following a function *λ*(*t*) according to the formula *N* (*t*) = *N*_0_*λ*(*t*), the distribution of *T*_*k*_ can be computed by

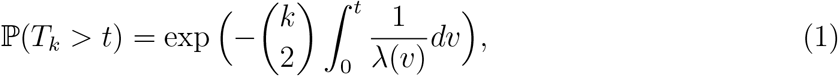

where 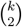 is the binomial coefficient 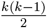.

The function *λ*(*t*) can be recovered in terms of the distribution of *T*_*k*_, by taking the log and then the derivative as in [50]. We have then

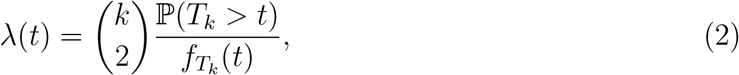

With 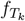, the pdf of *T*_*k*_.

We can note from equation 2 that the identifiability problem pointed out by [50] for the *T*_2_ distribution is also present for *T*_*k*_. That is, for any model of population evolution (structured population, isolation with migration, split models, admixture, etc.) and for a particular sampling configuration there will always be a model of population size changes under panmixia (given by equation 2) having exactly the same distribution of *T*_*k*_. Increasing *k* doesn’t solve the problem of spurious signals of population size change when the population is structured if only one *T*_*k*_ and one sampling scheme is used. Nevertheless, this identifiability issue can theoretically disappear (at least for the *n*-island model) when considering the joint information of more than one coalescence time. Indeed, [29] has shown for the *n*-island model, that it is possible if *T*_2_ and *T*_3_ are jointly used for a sample of size *k* = 3 haploid genomes.

Following [50] we can then define the IICR_*k*_ (Inverse Instantaneous Coalescence Rate for *k* genes) as follows

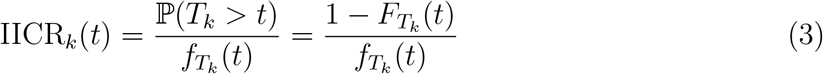

where 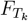 and 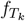 are the cumulative distribution function (cdf) and pdf of *T*_*k*_, respectively. From equation 2 we can see that, under panmixia, 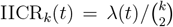 and will thus be proportional to the history of population size change (the *λ* function).

When the hypothesis of panmixia is not verified, the IICR_2_ cannot be interpreted as representing *λ*(*t*) [50]. It is thus reasonable to hypothesize that the same is true of 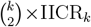 for *k >* 2 even though the properties of the IICR_*k*_ as a function of the demographic model, sample size and sampling scheme are not known and need to be clarified. The 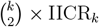 can be seen as a summary statistic [16] and understanding how it varies with the sampling scheme and size can inform us on the demographic models that can and cannot explain our data. For instance, in the case of a 2D stepping-stone of size 3 *×* 3, there are 11 different ways of taking a sample of two haploid genomes, each of them might give us a different curve for the corresponding IICR_2_. When increasing the sample size, the number of possibilities for the sample configuration increases considerably: we can count 42 different ways of sampling *k* = 10 haploid genomes in a symmetrical *n*-island model with 10 demes, and as many as 42 [1] different IICR_10_ for the same model just by changing the sampling vector.

We thus introduce the notation IICR_*k,v*_ where *v* is a sampling vector indicating how many haploid genomes we take from each deme in a given structured population model. We will use the same convention for the coalescence times, which will be denoted *T*_*k,v*_.

### 2.2 Exact computation of the IICR_*k*_

The different IICR_*k,v*_s can be derived analytically for any model for which the distribution of coalescence times, *T*_*k,v*_, is known, by equation (3). For most structured models, no closed expression is known, but the IICR_*k,v*_ can be numerically computed using the transition matrix approach of [33], as it has been done in [61] for the IICR_2_. This requires to construct the transition matrices of the demographic models of interest for the desired values of *k*.

Let us consider a model with *n* populations (or demes) numbered from 1 to *n*, with *s*_1_, …, *s*_*n*_ their respective sizes, 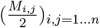 the normalized backward migration rates from deme *i* to deme *j* and 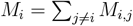.Let us sample *k* (*k* ≥ 2) genes from the population at the present (*t* = 0), the sampling vector can be denoted by *v* = (*v*_*i*_)_*i*=1…*n*_ where *v*_*i*_ is the number of genes in deme *i*. Note that 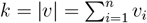.In order to trace the lineages back to the first coalescent event, we only need to consider the set *E*_*n,k*_ = {*v* ∈ ℕ ^*n*^, *k* − 1 ≤ |*v*| ≤ *k*}, since |*v*| = *k* − 1 means that two of the *k* lineages have coalesced (see [61] for the whole states space and details about migration rates).

The structured coalescent is then the Markov chain evolving in the following way. It can change from one state *v* ∈ *E*_*n,k*_ to another state *w* ∈ *E*_*n,k*_ either by a migration event (which implies that |*w*| = |*v*|) or by a coalescence event inside a deme (which implies that |*w*| = |*v*| − 1). The corresponding transition rate matrix can be constructed as:

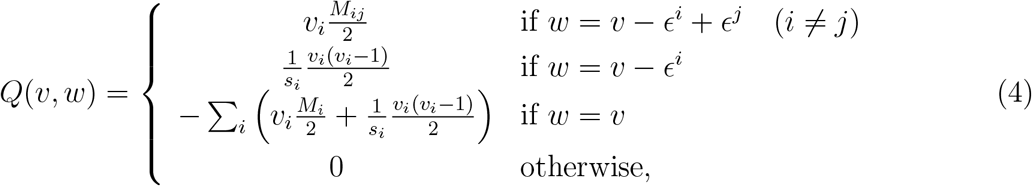

where *ϵ*^*i*^ is the vector whose components are 1 on the *i*^th^ position and 0 elsewhere. The matrix *Q* describes two types of possible events for each configuration *v*:

- *w* = *v* − *ϵ*^*i*^ + *ϵ*^*j*^ when one lineage migrates (backward in time) from island *i* to island *j*. The rate of this migration is *M*_*ij*_*/*2 (migration rate to deme *j* for each lineage in deme *i*) times *v*_*i*_, the number of lineages present in deme *i*.
- *w* = *v* − *ϵ*^*i*^ denotes a coalescence event between two lineages in deme *i*, which reduces the number of lineages by one in this deme. This occurs only if *v*_*i*_ ≥ 2. If this is not the case we can see that *v*_*i*_(*v*_*i*_ − 1) = 0. The term *v*_*i*_(*v*_*i*_ − 1)*/*2 is the number of possible pairs among the *v*_*i*_ lineages. This term is multiplied by 1*/s*_*i*_ since the *i*^th^ island has a normalized population size equal to *s*_*i*_, and 1*/s*_*i*_ is then the coalescence rate for each pair of lineages in this island.

Let us take the example of a population of 2 demes and *k* = 3. *E*_2,3_ is then a set of 7 elements: *E*_2,3_ = {(3, 0), (2, 1), (1, 2), (0, 3), (2, 0), (1, 1), (0, 2)} where we have chosen an arbitrary order of the elements. With this order, the transition rate matrix is:

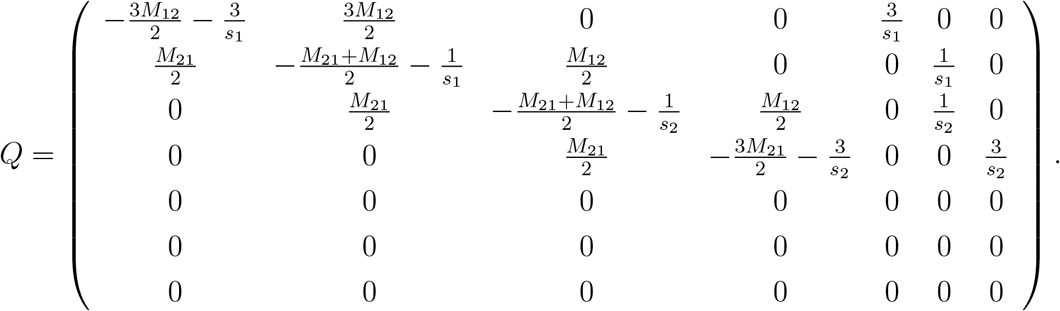

The last three states are absorbing because they correspond to the states when coalescence occurred. Let us note that we could also use a smaller matrix by merging these three states into a single “coalescent state”. This is acceptable if we are only interested in the first coalescence (here *T*_3_). Making them distinguishable states allows us to keep track of where the last two lineages are, so we could be able to compute the distribution of the coalescence time of the remaining two alleles.

Once the transition matrix is defined, Markov chain theory provides the tools allowing to compute the distribution of *T*_*k,v*_ with the matrix exponential

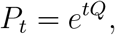

where the matrix coefficient *P*_*t*_(*v, w*) is the probability that the Markov process is in the state *w* at time *t* knowing that it started from the state *v* at time *t* = 0. We then obtain the cumulative distribution function (cdf)

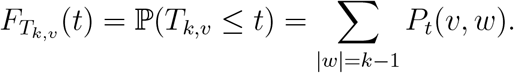

The pdf of *T*_*k,v*_ is by definition the derivative of 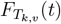, it can thus be computed from the matrix *P*_*t*_ by using the property

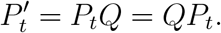

We can thus write

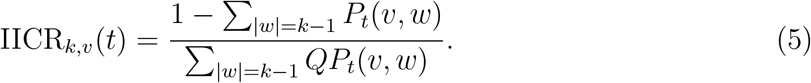

Details of those derivations can again be found in [61].

Taking back our example, if we sampled three genes by taking two in the first island and one in the second, the Markov chain would initially be in the state 2, and since the coalescence states are the numbers 5, 6 and 7, the IICR_3,(2,1)_ would be :

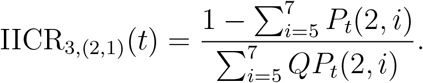

### 2.3 Estimation of the IICR_*k*_

With increasing *k* values and number of demes, the number of possible states and the size of the matrices quickly increase, making this approach computationally prohibitive. When this is the case, the IICR_*k,v*_ can still be estimated for any demographic model for which *T*_*k,v*_ values can be simulated, by computing

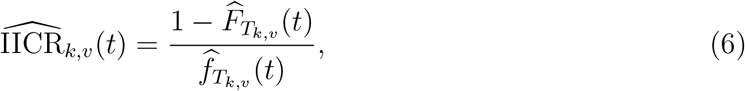

where 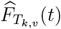 is the empirical *cdf* of *T*_*k,v*_ evaluated at some point *t* and 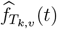 is an estimation of its density around *t*.

For each scenario and sample configuration presented below we simulated 10^6^ independent *T*_*k,v*_ values. With real or simulated genomic data it is also possible to estimate the 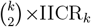. For *k* = 2 this can be done with the PSMC and MSMC methods of [44] and [65]. For larger *k* values this can only be done with the MSMC method of [65].

In the following section, we explore the 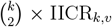 under several structured models commonly used in population genetics. We explore how 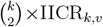 curves change under the same model when *k* and *v* vary. We are also interested in comparing how the three approaches above, namely the simulation of *T*_*k,v*_ values, the matrix approach, and the application of the PSMC and MSMC methods to simulated data, provide similar results. We also tested the MSMC2 method as it has been presented as an improvement over the latter.

The R and Python scripts allowing to construct the transition rates matrix for the general case and to compute and plots the 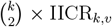 based on formula (6) are freely available here https://github.com/willyrv/IICREstimator/tree/master

## 3 Results

### 3.1 The shapes of 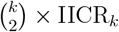 curves under structured models

Figure 1 shows the results obtained for the 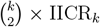 assuming an *n*-island model (*n* is the number of islands), where we varied the sample size (*k*) and migration rate (*M* = *M*_*i*_ for all *i* ≤ *n*), but keeping the sampling scheme constant, with all samples obtained from the same deme. This figure shows that, as *k* increases, the 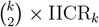 is shifted left (towards recent times) and downward (towards smaller values) compared to the IICR_2_, except in the recent time where all curves converge towards the size of a deme, as expected [50]. This shifting effect decreases when migration increases, as expected, since the *n*-island model tends towards a panmictic model where all 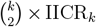 are expected to be equal to the total size, *n × N*. This effect is observed for both the exact 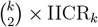 and for the approximate 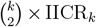 obtained through simulations (continuous *versus* discontinuous stepwise curves, respectively, see Figure S1 for additional *M* values). Note also that comparing panel a and b we also see that as *M* increases the 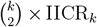 is shifted downward and leftward to more recent times as shown previously by [50] for the IICR_2_. Thus, increasing *k* for a constant *M* appears here to have a *qualitatively* similar effect to increasing *M* for a given *k*.

**Figure 1:**
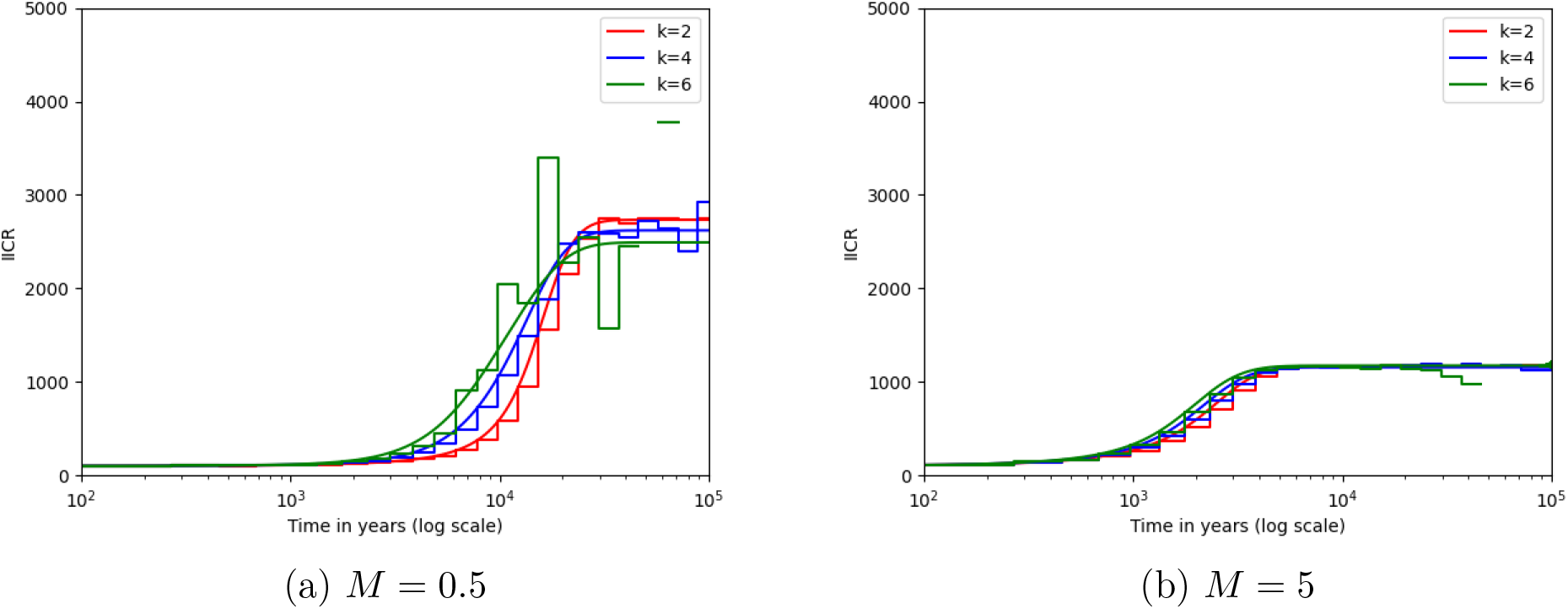
Sample size effect on the 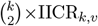. The 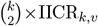 were obtained for a *n*-island with *n* = 10 demes. The two panels correspond to models with different migration rates, *M* = 0.5 (panel a) and *M* = 5 (panel b). The *k* haploid genomes were sampled in the same deme (*v* = (*k*, 0, …, 0)). Exact computation (formula (5)) are plotted as solid curves, and estimated 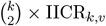 (formula (6)) are plotted as stepwise curves. For each value of *k*, the `ms` command used here was `ms k 100000 -I 10 k 0 0 0 0 0 0 0 0 0 M -T -L`, where *k* and *M* are replaced by the values above. The IICR values are scaled taking into account that *M* = 4*mN*_0_ where *N*_0_ = 100 individuals by deme. The time values are scaled assuming a generation time *g* = 25 years and a mutation rate *µ* = 1.25 *×* 10^−8^.

Figure 2 shows the effect of increasing *k* while sampling diploid individuals in different demes rather than in the same deme. As in Figure 1, an *n*-island model was assumed and different migration rates are represented in the different panels. We observe again a shift of the 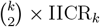 as *k* increases. The shift however differs from the effect observed in Figure 1. Here the 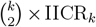 curves differ significantly in the recent past where the 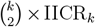 is not equal to the deme size except for *k* = 2. Another important difference is that even when migration is high, the curves maintain a vertical shift in the recent past. In other words, the sampling scheme creates a “population size effect” in the recent past that has little to do with any real size change, or any variation in deme size across the metapopulation. See Figure S2 for additional *M* values.

**Figure 2:**
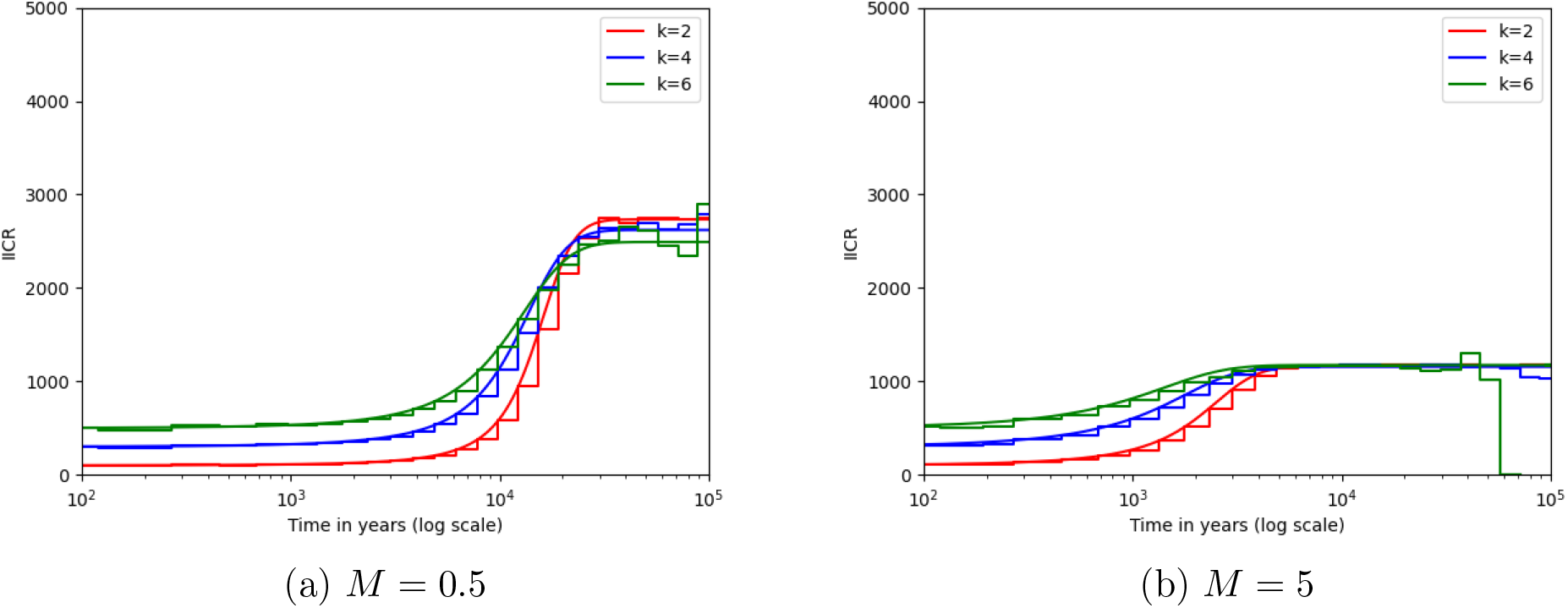
Sample size and sampling scheme effect on the 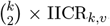. The 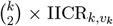’s are obtained and plotted as in Figure 1, the only difference is the way to sample: when *k* = 2, both individuals are in the same deme, when *k* = 4 we take 2 on one deme and 2 in an other, and when *k* = 6, we take three pairs of individuals in three different demes. The `ms` command are then respectively `ms 2 100000 -I 10 2 0 0 0 0 0 0 0 0 0 M -T -L`, `ms 4 100000 -I 10 2 2 0 0 0 0 0 0 0 0 M -T -L` and `ms 6 100000 -I 10 2 2 2 0 0 0 0 0 0 0 M -T -L`.

In Figure 3 we now explore, for a fixed value of *k* = 4, the effect of the sampling scheme in a stepping-stone model (See Figure S3 for the different sampling schemes). Interestingly, this example shows again a new behaviour. These examples show that even if *k* is the same, and the recent and ancient plateaux of the 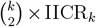 are the same, the temporal dynamics differs when the samples are obtained from different demes and analysed together. This can even generate a hump in sampling schemes 6 and 7. A similar hump observed in a MSMC curve would be interpreted as non monotonous change in size. As in Figure 2, increasing migration does not lead to an overlap of the 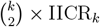 curves.

**Figure 3:**
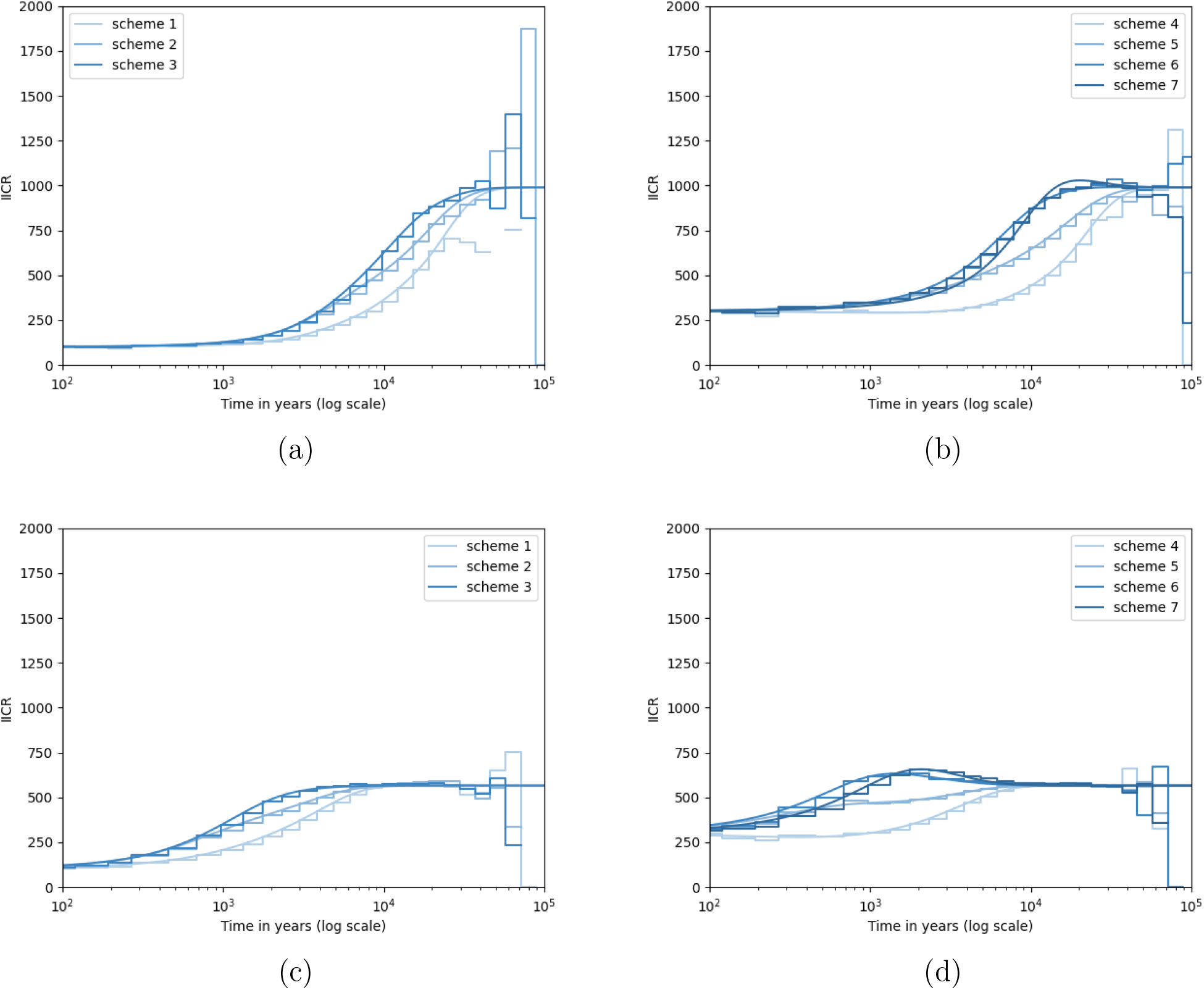
Sampling scheme effect on the 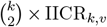 in a linear stepping-stone: sampling scheme effect. The 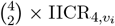 curves were obtained for a linear stepping stone of 5 islands, with the migration rates *M* = 0.5 (panels (a) and (c)) and *M* = 5 (panels (b) and (d)). Note that here the migration rate *M* = *M*_*ij*_, constant for all *i, j* ≤ 5). We only considered sampling schemes *v*_*i*_ with *k* = 4 haploid individuals. For the first three sampling schemes (noted *scheme i* in the legend, for *i* = 1, 2 and 3) we sampled the 4 genomes (i.e. two diploid individuals) in the same deme, with scheme *i* corresponding to deme number *i* (i.e. in scheme 1 all individuals are sampled in the first deme of the linear stepping stone). For schemes *i* = 4, …, 7 we sampled one diploid individual in deme 1 and the other in deme 2, 3, 4 or 5, respectively (see Figure S3). Exact computation (formula (5)) are plotted by solid curves, and estimated 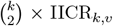 (formula (6)) are plotted by stepwise curves. For each value of *M*, and for sampling scheme 1, the `ms` commands used here is `ms 4 1 -I 5 4 0 0 0 0 0 -m 1 2 M -m 2 3 M -m 3 4 M -m 4 5 M -m 5 4 M -m 4 3 M` `-m 3 2 M -m 2 1 M -T`. The IICR and times values are scaled like in Figures 1 and 2.

Thus, figures 2 and 3 show that even under scenarios with high *M* values the sampling scheme can create differences in observed PSMC or MSMC curves which would be interpreted as representing different demographic histories in terms of change in *N*_*e*_.

Altogether these three figures show that significant differences between 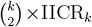 curves are expected when *k* changes for structured models, and that the curves can also change when *k* is fixed as a function of the sampling scheme, even for high migration rates. In the next figures we test whether the MSMC method is consistent with our theoretical results by simulating genomic data under the structured models and sampling schemes above.

### 3.2 The shapes of PSMC, MSMC and MSMC2 curves under structured models

Figure 4 shows the theoretical 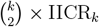 curves of the same *n*-island models represented in Figure 1 but with the corresponding MSMC curves. We can observe several patterns. First, the MSMC curves for *k* = 2, 4 and 6 start to follow the expected and corresponding 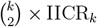 curves, at least in the most recent past. This suggests that the MSMC method does infer the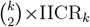, similar to the expectation under panmixia [66]. Second, the MSMC curves tend to infer humps or changes which are not predicted by the theory presented here. Such changes would be interpreted as changes in population size and may be due to stochastic variability as a consequence of imperfect information in the sequence data. Third, the fit to the expected curve gets worse as *k* increases. Finally, the MSMC curves seem to improve a little by becoming closer to each other when migration increases but this is not a major improvement when compared to the expected 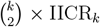 curve, since the MSMC curves are still quite different from the expected 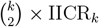curve.

**Figure 4:**
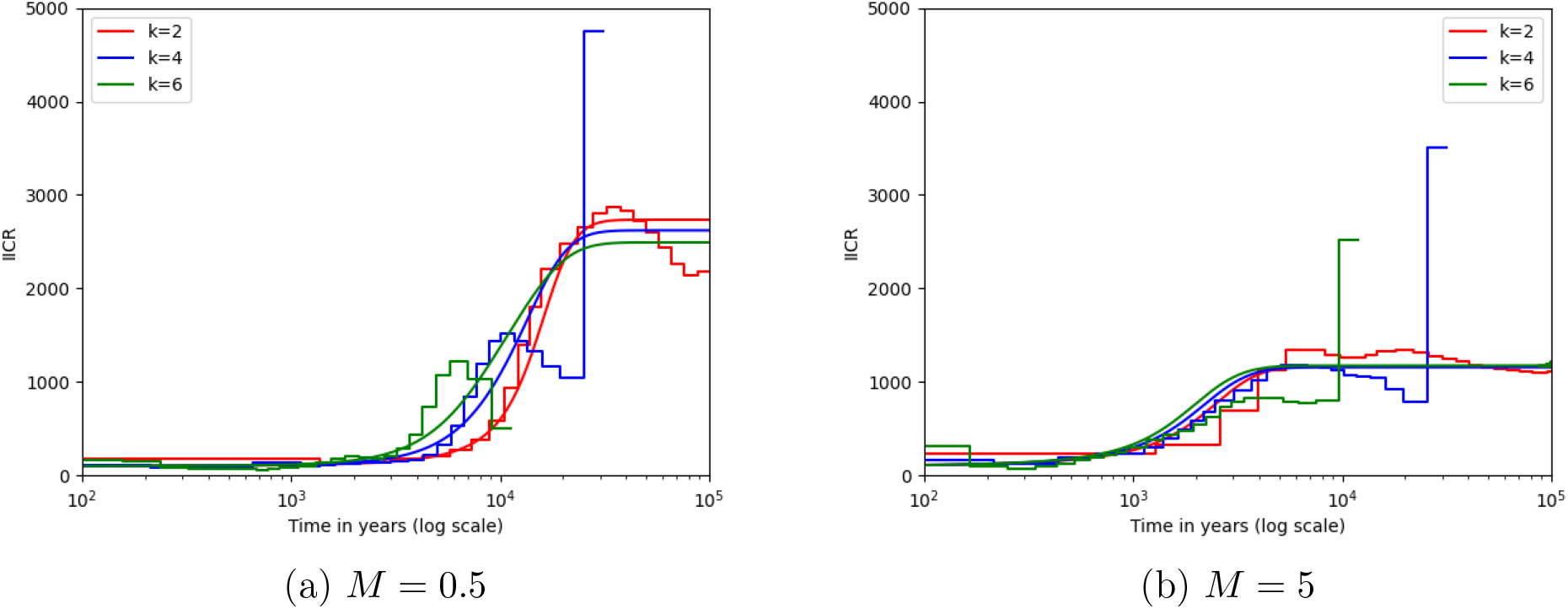
Sample size effect on MSMC plots and on the 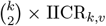. The 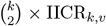 ‘s were obtained for the same models and sampling schemes as Figure 1. Here the stepwise curves are MSMC outputs using genomic data simulated with `msprime` in the following way: each sequence is composed by 10 chromosomes of length *l* = 3 *×* 10^8^ bp, the mutation rate is *µ* = 1.25 *×* 10^−8^ per bp per generation, the recombination rate *ρ* = 10^−8^ per bp per generation, and the maximum number of recombinational segments set to 8. The `msprime` commands are then `mspms k 1 -t 1500 -r 1200 300000000 -I 10 k 0 0 0 0 0 0 0 0 0 M -p 8 -T` with the respective values of *k* and *M*. Note that the `-t` and `-r` values are respectively obtained from the products 4*N*_0_*µl* and 4*N*_0_*ρl*.

Figures 5 and 6 compare the MSMC results for the same models and sampling schemes as in Figures 2 and 3, respectively. Again, these results suggest that the MSMC method infers the 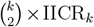 but suffers from a stochastic behavior which might be misleading when analysing real data. It is interesting to see that the MSMC curves do reproduce the humps of the sampling schemes 6 and 7 for the stepping stone, even if in an exaggerated way and at a different time.

**Figure 5:**
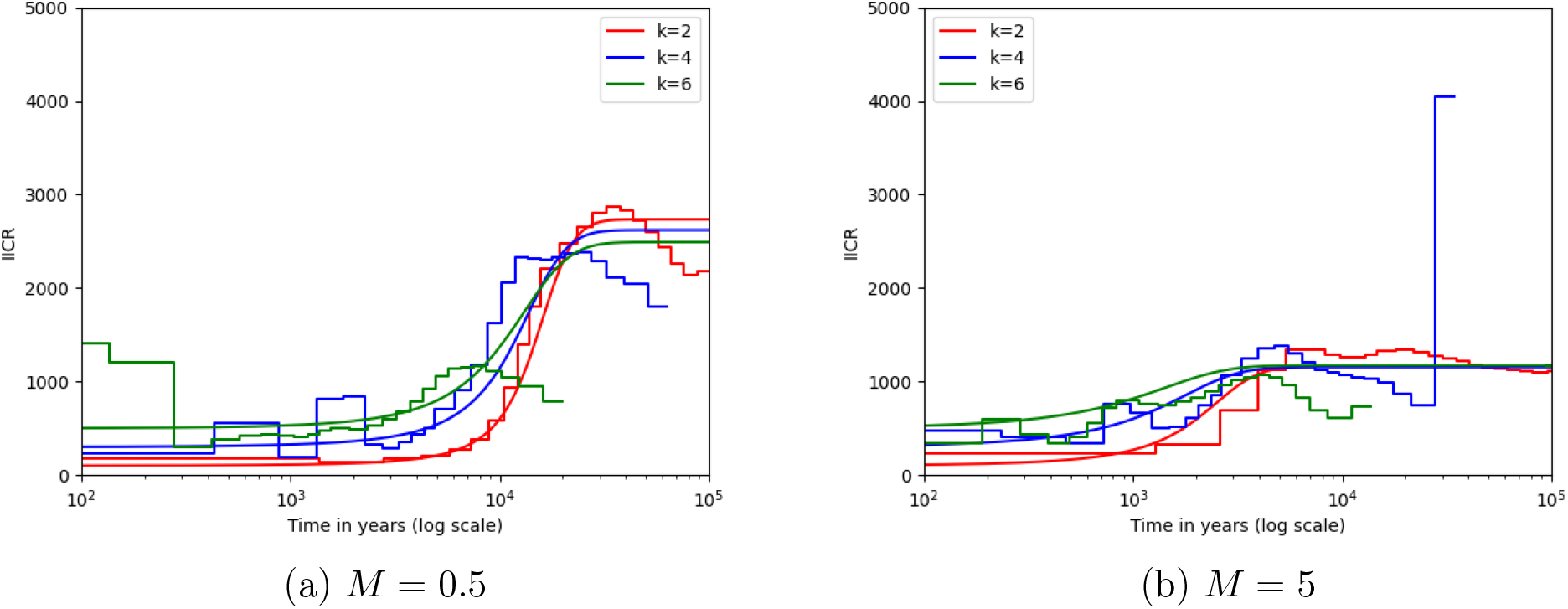
Sample size and sampling scheme effect on MSMC plots and 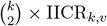.See Figure 2 for details regarding the model and sampling scheme.

**Figure 6:**
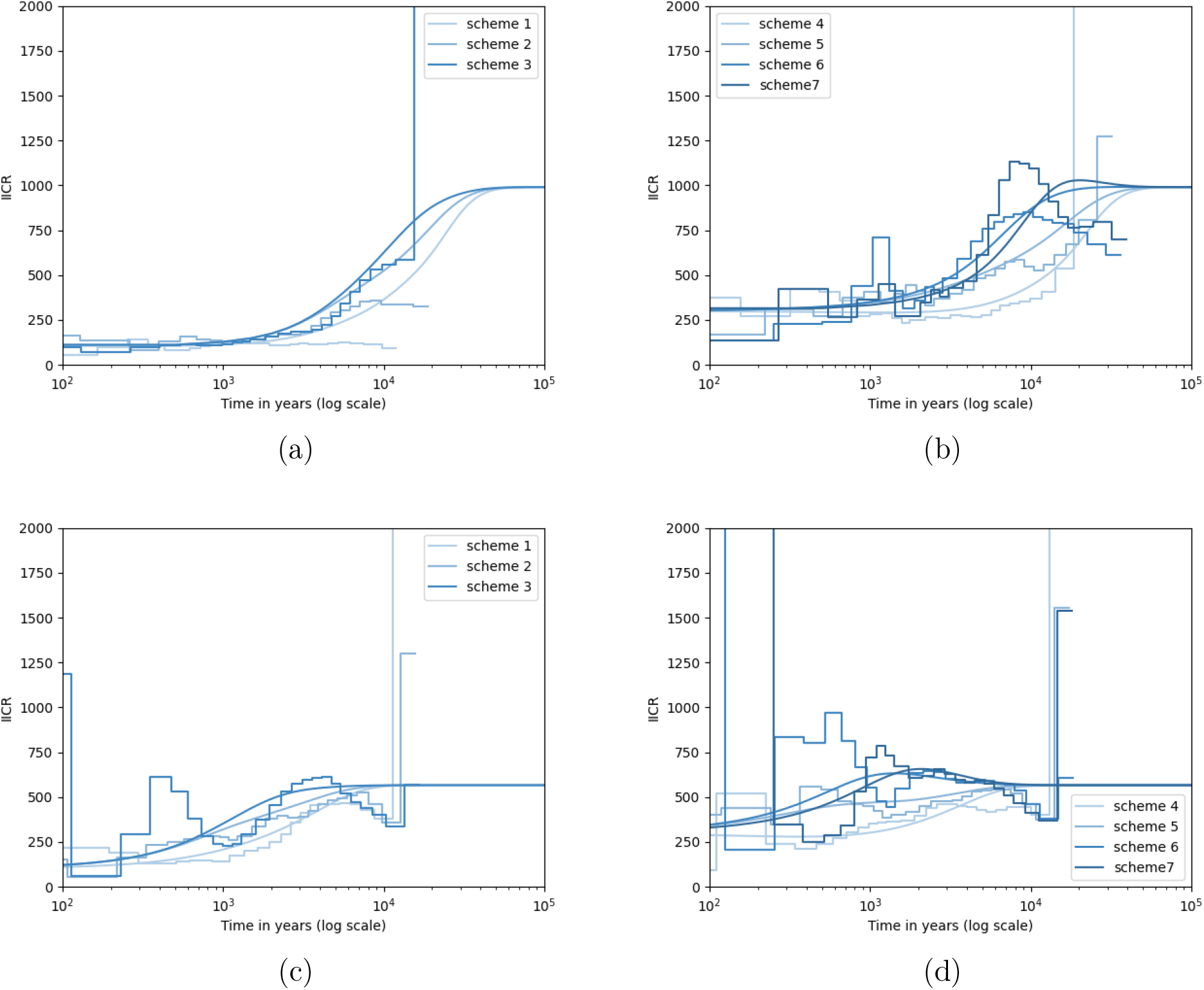
Sampling scheme effect on MSMC plots and 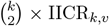 in a linear stepping-stone. See Figure 3 for the simulation details. The main difference with this figure is that we present the results for two *M* values, namely *M* = 0.5 (panels (a) and (c)) and *M* = 5 (panels (b) and (d)). The `msprime` commands for the simulation are `mspms 4 1 -t 1500 -r 1200 300000000 -I 5 4 0 0 0 0 0 -m 1 2 M -m 2 3 M -m 3 4 M -m 4 5 M -m 5 4 M -m 4 3 M -m 3 2 M -m 2 1 M -p 8 -T` for sampling scheme 1.

Altogether, these figures suggest that the MSMC method infers the 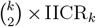 but that it may suffer from too much variability to be used as summary statistic for demographic inference, at least under the conditions used here. To test whether we could reduce the stochasticity issue mentioned above we ran ten independent runs for the first panel of Figure 5 and plotted both the independent runs and the average MSMC curves. While we found that the fit to the expected curve improved, there still were differences, particularly for *k* = 6 as can be seen in Figure S5 and S6

Given that the MSMC2 method was introduced as an improvement on the PSMC and MSMC methods [48, 66] we also applied the MSMC2 method to genomic data generated under several scenarios and sampling schemes. In Figure 7 we compare the PSMC, MSMC and MSMC2 curves obtained for a diploid genome (*k* = 2) sampled in one deme of an *n*-island model with different migration rates. The figure shows that the three methods seem to estimate or approximate the theoretical IICR_2_ curve, with different levels of stochasticity. The PSMC appears to produce the best results, followed by MSMC2, whereas MSMC continues to produce a hump that is not theoretically expected.

**Figure 7:**
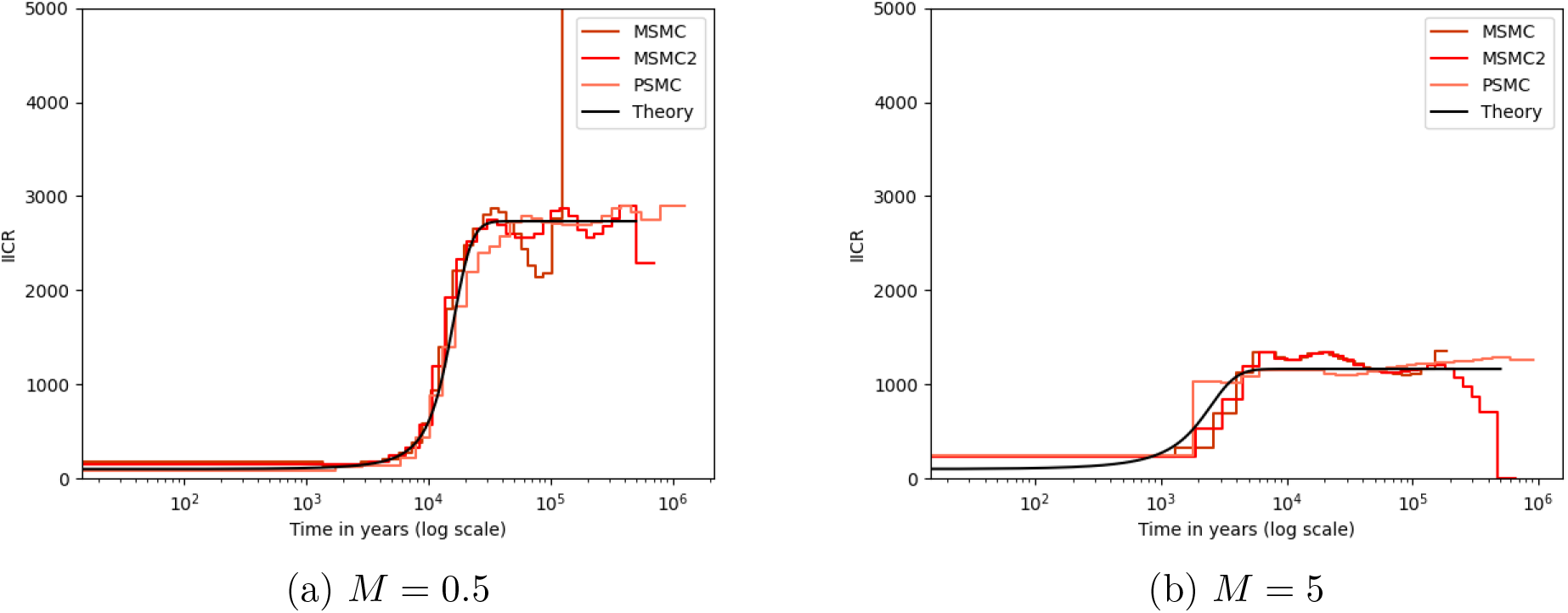
Sampling from the same deme: comparison of PSMC, MSMC and MSMC2 plots with the 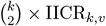. Here we focus on the IICR_2_ for the same *n*-island as in Figures 1 and 4, *n* = 10 and *M* = 0.5 (panel (a)) or *M* = 5 (panel (b)). We thus assume that *k* = 2, and that the two haploid genomes are sampled in the same deme. The theoretical black solid curve is computed with formula (5), and the PSMC, MSMC and MSMC2 outputs are obtained with the sequence simulated by the `msprime` command `mspms 2 10 -t 1500 -r 1200 300000000 -I 10 2 0 0 0 0 0 0 0 0 0 10 -p 8 -T`.

In Figure 8 we plot the results of MSMC2 for the same sampling schemes as in Figure 3. Compared to the results obtained with the MSMC method the figures show much less stochasticity, as expected given that the method appears to use all pairwise comparisons among haploid genomes [48, 66], but the stepwise curves differ significantly from the expected 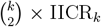 curves (see Figures S7, S8 and S10 for additional inferences and tests). The MSMC2 method thus differs from the PSMC and MSMC methods in that it does not infer the 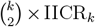, but it clearly uses information from more than two haploid genomes. When the genomes are sampled in the same deme this means that the curve obtained is possibly the IICR_2_. If correct the MSMC2 method would provide a better estimate of the IICR_2_ than the PSMC and MSMC methods. However, when the genomes are sampled in different demes, the MSMC2 method produces smoother curves but these curves should not be compared to MSMC curves because they do not estimate the same object (outside panmixia, at least). Also, all our simulations were done with no population size change, which thus suggests that these curves do not infer population size change under the structured models simulated here.

**Figure 8:**
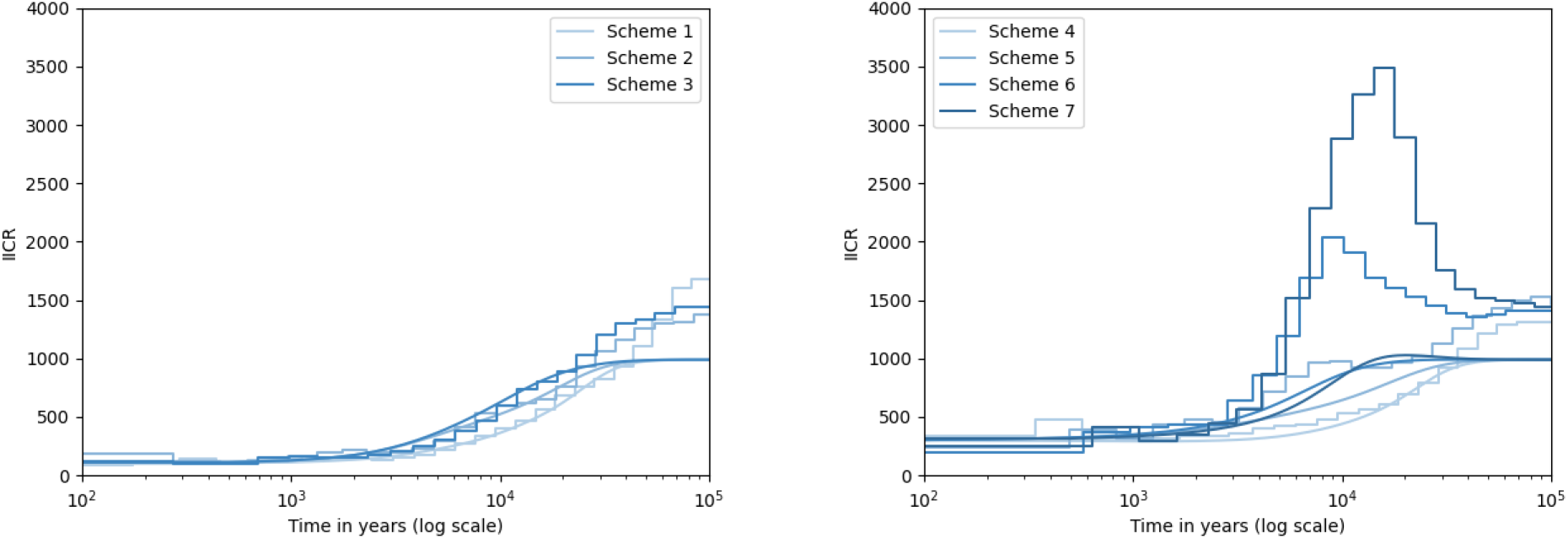
Sampling from different demes in a linear stepping-stone: sampling scheme effect on MSMC2 plots (stepwise curves) and 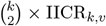 (continuous curves) for *k* = 4. The theoretical 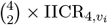 curves were obtained for the linear stepping stone with *n* = 5 demes, *M* = 0.5, and the sampling scheme *v*_*i*_ as in Figure 3. The stepwise curves are the outputs of MSMC2 for the simulated sequence with the same `msprime` commands as Figure 6.

## 4 Discussion

### 4.1 Phylogeography and the limits of “demographic history”

Phylogeographical studies have shown for decades that the distributions of most animal and plant species have changed in their recent past [35, 34, 84, 67, 3, 2, 64, 78]. This suggests that the habitats of most species have gone through periods of contraction, fragmentation and expansion. The populations of these species have thus been partly isolated and brought into contact as a consequence of environmental changes [34] and have thus gone through periods of increased or reduced gene flow, with changes in population sizes. How changes in population size and connectivity have interacted and how much each of these may have been more important for different species is difficult to say today. To answer these questions the properties of genetic data under alternative scenarios need to be better understood [2]. This may help us identify the statistics that can inform us on such changes, and perhaps clarify what the statistics we compute represent. This would be similar to the work previous researchers who studied the properties of Tajima’s D and other statistics during the 1990s and 2000s [73**? ?** ].

The phrase “demographic history” is often meant to represent a history of population size changes. While this is not a problem *per se*, an increasing number of studies suggest that, as soon as populations are structured, the changes in size inferred by several approaches beyond those analysed here may be significantly misleading [79]. Calling “demographic history” a series of inferred population size changes that may never have happened is questionable [17, 16**?** ]. What population geneticists reconstruct as “demographic trajectories” are actually curves whose meaning is often unclear and inconsistent across methods and studies [10]. For instance, the PSMC, MSMC and MSMC2 curves are not expected to represent the same objects under structured models and none of them is unambiguously representing population size changes. Still, many studies use them interchangeably and with other methods to infer and tell a history of changes in *N*_*e*_ (e.g. [11]).

### 4.2 What the IICR_*k*_ tells us about the PSMC, MSMC and MSMC2 curves and “demographic history”

In the present study we extended the work of [50] and confirmed that the PSMC method estimates the IICR_2_. We also extended the matrix approach of [61] to the 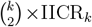 making it possible to compute the 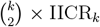 efficiently. Additionally, we found that the MSMC method estimates the 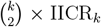 for *k* ≥ 2, even though the stochasticity of the MSMC results makes it difficult to fully validate this statement. We thus warn users regarding the results obtained from the MSMC method with real data. One additional and important result is that there is no particular reason to expect these two methods to produce the same curves when population structure is present and significant and when sample size or sample configuration change. For instance, under structure, if we apply the PSMC method to three diploid individuals we will obtain three PSMC curves, which will differ from the MSMC curves that we will obtain by analysing these individuals together or in pairs. This effect will be stronger if the sampling varies (when individuals come from different demes) but differences between PSMC and MSMC results were observed when the individuals were sampled in the same deme. This can be seen when we sample an increasing number of individuals from the same deme under a simple *n*-island (Figure 1) or stepping stone (Figure 3). These two methods cannot be seen as representing the “demographic history of the population” whichever meaning one might want to give to that phrase. Our results also contribute to explaining the results of [10] who were among the first to clearly state that the MSMC, PSMC and SMC++ methods produced inconsistent results.

The MSMC2 method was introduced as an improvement over the PSMC and MSMC methods [48, 66]. However, we found that MSMC2 produces additional curves that do not correspond to the expected 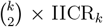 in the general case, even though they may approximate the IICR_2_ in the recent past when all individuals are sampled in the same deme. For instance, in the first panel of Figure 8 we found that when two diploid individuals were sampled in the same deme in a stepping stone, the MSMC2 curves were following the IICR_4_ in the recent past but tended to significantly overestimate the plateau in the ancient past. Under more complex sampling schemes (second panel of Figure 8) the MSMC2 curves continue to significantly overestimate the ancient plateau but they also exhibit major changes in the y-axis that exaggerate very moderate humps expected in the theoretical 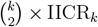. With real data such curves would suggest a history with an ancient large population that went through a major population growth and then crashed around 10,000 **years** ago. This is misleading given that all the models studied here were stationary without any change in size or connectivity. More work is thus necessary to clarify the theoretical underpinnings of the MSMC2 and other published methods that produce similar curves and use different types of information, such as the SMC++ method developed by [76].

We focused here on methods that require small sample sizes that are attainable for non-model and endangered species, and are increasingly being used for endangered species [83, 74]. The problem discussed here is however general and likely affects many methods based on the standard (non structured) coalescent. The PSMC, MSMC and MSMC2 methods have been introduced and interpreted mainly as representing changes in *N*_*e*_. The reason for this is that under the standard coalescent the coalescence rate is inversely related to *N*_*e*_. The implicit assumption is that population structure, if it exists, is limited and negligible and the inferred changes in *N*_*e*_ should be reasonably close to actual changes, that themselves reflect changes in census size. While this may seem reasonable at first, an increasing number of studies have demonstrated that current methods, when tested, are not robust to population structure [79, 6, 56, 72, 17, 32, 55, 50]. So much so that as soon as population structure becomes relevant the relationship between inferred changes in *N*_*e*_ and real changes in *N*_*e*_ under structure can change from a positive to a negative correlation through no correlation at all. Thus, one can infer a bottleneck when a population is actually growing or stationary [79, 17, 50]. Similarly, one can infer an expansion when the population size is decreasing or stationary [50].

The effect of population structure is not limited to the inference of non-existing population size changes. Several studies have now shown that spurious admixture events may also be inferred [19, 20, 77] and it is likely that the detection of selection in the genome might also be influenced by population structure and gene flow [40, 12, 39, 68]. Ignoring population structure will thus lead to the detection and quantification of events that may never have taken place or may have taken place at different times from those inferred. Since much of what we understand of the recent evolutionary history of species is based on our interpretation of these curves or on other patterns observed in genomic data, this may lead to serious misunderstandings and misrepresentations of the evolutionary history of species.

### 4.3 Can the IICR_*k*_ help improve demographic inference and model choice

In an insightful study, [10] noted that the PSMC curves obtained from human genomes were different from the curves obtained using the MSMC method with 4 or 8 haploid genomes, which were themselves different from each other. They additionally noted that these curves were also very different from those obtained using the SMC++ method of [76] which uses additional information from the SFS (site frequency spectrum). And all these curves differed from the curves of size change obtained when one simulated genomes under scenarios inferred using the dadi method [31], which only uses the SFS as a summary statistics. They also noted that no inferred history could reproduce the LD patterns observed in humans, whereas three methods could reproduce the average heterozygosity along the genome. The authors interpreted this as a suggestion that the PSMC and MSMC methods were somehow biased. Our results suggest another interpretation where the different methods were not necessarily producing wrong results. Rather, it is the interpretation of the curves that is problematic as this interpretation changes when one accounts or not for population structure. The genomic data analysed in their study were either simulated under the tree population model of [31] or using panmictic models with population size changes inferred by the different methods. Under a panmictic model the PSMC, MSMC and MSMC2 methods should all produce the same results with differences only related to precision in the recent or ancient past. Under structured models sampling becomes important and the different methods are expected to generate different curves. And none of these curves should be used to represent a poorly defined “demographic history”. Rather, these curves should be used as complex summary statistics for demographic inference [16, 77].

In a recent study, [4] developed a method which uses the PSMC curve as a summary statistic and infers the parameters of a non-stationary *n*-island model. This method, called SNIF for Structured Non-stationary Inference Framework, aims at identifying periods during which connectivity (gene flow) changed under the assumption that the number of islands and their sizes are constant (https://github.com/arredondos/snif). This method has been validated on a large parameter space and used to analyse human genomes. The authors found that the model inferred by [31] was unlikely to represent a reasonable model to explain human PSMC plots. This suggested that current models of human evolution assuming a tree structure with three main “continental” populations may significantly misrepresent recent human evolutionary history.

One important limitation of SNIF is that it assumes an *n*-island model, which is not realistic for many species. For instance, humans exhibit clear isolation by distance patterns that suggests that spatial structure should be incorporated [19, 77]. Future approaches should thus integrate space, using for instance stepping-stone models [21], as a first step to identify structured models that can explain PSMC, MSMC, MSMC2 and SFS plots altogether.

### 4.4 The IICR_*k*_ and a possible future of “demographic inference”

In principle, models used to represent the demographic history of populations should be as simple as possible while at the same time incorporating as much as possible of the complexity of natural populations to make meaningful inferences in relation to the questions asked by researchers. A complex and realistic model could include changes in population size, in population structure, and in connectivity [25, 36]. It could also integrate geographical features and differences between males and females, such as sex-biased admixture or migration [59] or social structure and mating systems [54] and selection [40, 39]. However, the number of parameters required for such realistic models might be so large that the inference process would be difficult and perhaps questionable even with genomic data. As a consequence population geneticists have understandably focused on only some of these aspects of the demographic history of species [36]. However, the price to pay for this simplification is that spurious events may be inferred and discussed in heated debates, when they might actually have never taken place.

By expliciting how the IICR_*k*_ can be efficiently obtained for a very large family of demographic models the current study may contribute to clarifying concepts that are central to demographic inference. Also, by providing a framework to quickly explore a large parameter space this should make it easier to develop new simulation-based inferential methods. The IICR_*k*_ could be used in an approximate Bayesian computation framework [9, 8] or with some of the new simulation-based approaches that are being developed [18, 47].

There have been major advances in the way we can simulate genomes [71] and use new statistical approaches to either better explore complex parameter spaces or use smaller numbers of simulations efficiently [8, 45]. For instance, one fruitful strategy consists in identifying several plausible models integrating information from other fields such as archaeology, ecology and palaeoclimatology and in using model choice approaches to identify the best model or models and the best fitting parameters for these models. There is renewed interest for such approaches as they theoretically allow us to have a more objective approach to the stories we tell [9, 22, 63, 4, 77, 82].

## Acknowledgements

We are grateful to the IRP BEEG-B (International Research Project - Bioinformatics, Ecology, Evolution, Genomics and Behaviour) (CNRS, Université Paul Sabatier, cE3c and IGC) for facilitating travel and collaboration between Toulouse (EDB, IMT and INSA) and Lisbon (IGC and cE3c). This work was partly performed using HPC storage and computing resources from genotoul and CALMIP (Calcul Midi Pyrénées, Grant 2012 - projects 43, 44, and 2015, project) bioinformatics platforms from Toulouse, Midi-Pyrénées, France. LC’s and OM’s research was also supported by the 2015-2016 BiodivERsA COFUND call for research proposals, with national funders ANR (ANR-16-EBI3-0014) the Fundação para a Ciência e Tecnologia (reference: Biodiversa/0003/2015 and PT-DLR 01LC1617A). LC also received support from the FCT project ref. PTDC-BIA-EVL/30815/2017. LC, OM and SB additionally received support from the DevOCGen project, funded by the Occitanie Regional Council’s “Key challenges BiodivO” program. JC’s research was supported by the Interdisciplinary Thematic Institute IRMIA++, as part of the ITI 2021-2028 program of the University of Strasbourg, CNRS and Inserm, supported by IdEx Unistra (ANR-10-IDEX-0002), and by SFRI-STRAT’US project (ANR-20-SFRI-0012) under the framework of the French Investments for the Future Program. This work was also supported by the LABEX entitled TULIP (ANR-10-558 LABX-41 and ANR-11-IDEX-0002-02) as well as the IRP BEEG-B (International Research Project Bioinformatics, Ecology, Evolution, Genomics and Behaviour). We acknowledge an Investissement d’Avenir grant of the Agence Nationale de la Recherche (CEBA: ANR-10-LABX-25-01).

## 5 Supplementary Material for

### I. Introduction

As explained in the main text the IICR_*k*_ is the Inverse Instantaneous Coalescence Rate among *k* lineages, and is related to the first coalescent event. The generalization from two to *k >* 2 lineages requires a scaling factor such that it is 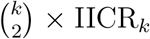 that is equivalent to *N*_*e*_ in a panmictic population. When *k* = 2 the scaling factor becomes 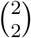 which is equal to one and explains why the original IICR is equal to *N*_*e*_. Throughout the main manuscript we computed the 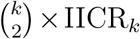 using three different approaches. The first requires a program able to simulate *T*_*k*_ values under the model and sampling schemes of interest. The second one uses a *Q*-matrix approach as in [**?** ]. Finally, we also simulated genomic data and provided the simulated data to the PSMC, MSMC and MSMC2 methods.

The Python and R scripts required for all the simulations and computations are based on the IICREstimator set of Python scripts (https://github.com/willyrv/IICREstimator) written by Willy Rodriguez for the [16] article and R scripts written by Cyriel Paris (https://github.com/aljounia/scripts_IICRk) to generate the theoretical 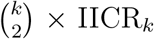 in his Masters thesis [Paris].

In addition all the Python and R scripts used to produce the curves, genomic data and to run the PSMC, MSMC and MSMC2 programs are available here: (see Annex II.).

The parameters used in this study are detailed in the main manuscript but can be summarized as follows:

- for the sample size we used the following *k* values: 2, 4, 6,
- for the number of islands we used *n* = 10 (for the n-island model) and *n* = 5 for the stepping stone models,
- several sampling scheme were used and are detailed below with the corresponding figures,
- we assumed that the size of all demes was the same and constant through time,
- we used several migration rates for the different models, namely *M* = 0.5, 1, 5, 10 with *M* = 4*N*_0_ *× m*. The migration matrices for the n-island model and for the stepping stone model are represented below (matrix A and B, respectively):

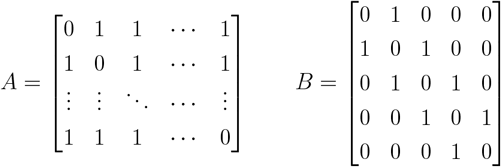

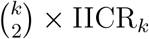 computation

For all the demographic scenarios for which we computed the IICR_*k*_ we did the following:

- we simulated 10^6^ *T*_*k*_ values
- we used Hudson’s *ms* program (the specific commands are given in the section II.)
- for the scaling of the 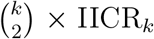 we assumed that the size of the population was *N*_0_ = 100 and the generation time was *g* = 25.

#### Simulation of genomes and the PSMC, MSMC and MSMC2 methods

The genomic data were simulated with msprime [43] since it allows to simulate genomes using the SMC approximation, which is assumed by the PSMC, MSMC and MSMC2 method. We used the following values for the simulations:

- for the recombination rate *ρ* =1.0 10^−8^ per bp per generation,
- for the mutation rate *µ* =1.25 10^−8^ per bp per generation,
- for the length of chromosomes 310^8^ bp (*i*.*e*. 300 MB),
- for the length of the genome 310^9^ (*i*.*e*. 3 GB composed of ten 300 MB-chromosomes),
- for the population size *N*_0_ = 100 and for the generation time *g* = 25.

### II. Theoretical IICR

The command in the parameter file necessary for IICREstimator (see appendix I.) was written as follow:

~~~
ms k 1000000 -I 10 k 0 0 0 0 0 0 0 0 0 M -T -L
~~~

with k, the sample size (k=2,4,6) and *M* = 4 *× m × N*_0_, the migration rate (M=0.5,1,5,10). In this example, 10^6^ *T*_*k*_ have been sampled from an n-island model with 10 islands and all the individuals were sampled in the same deme (Figure S1).

**Figure S1:**
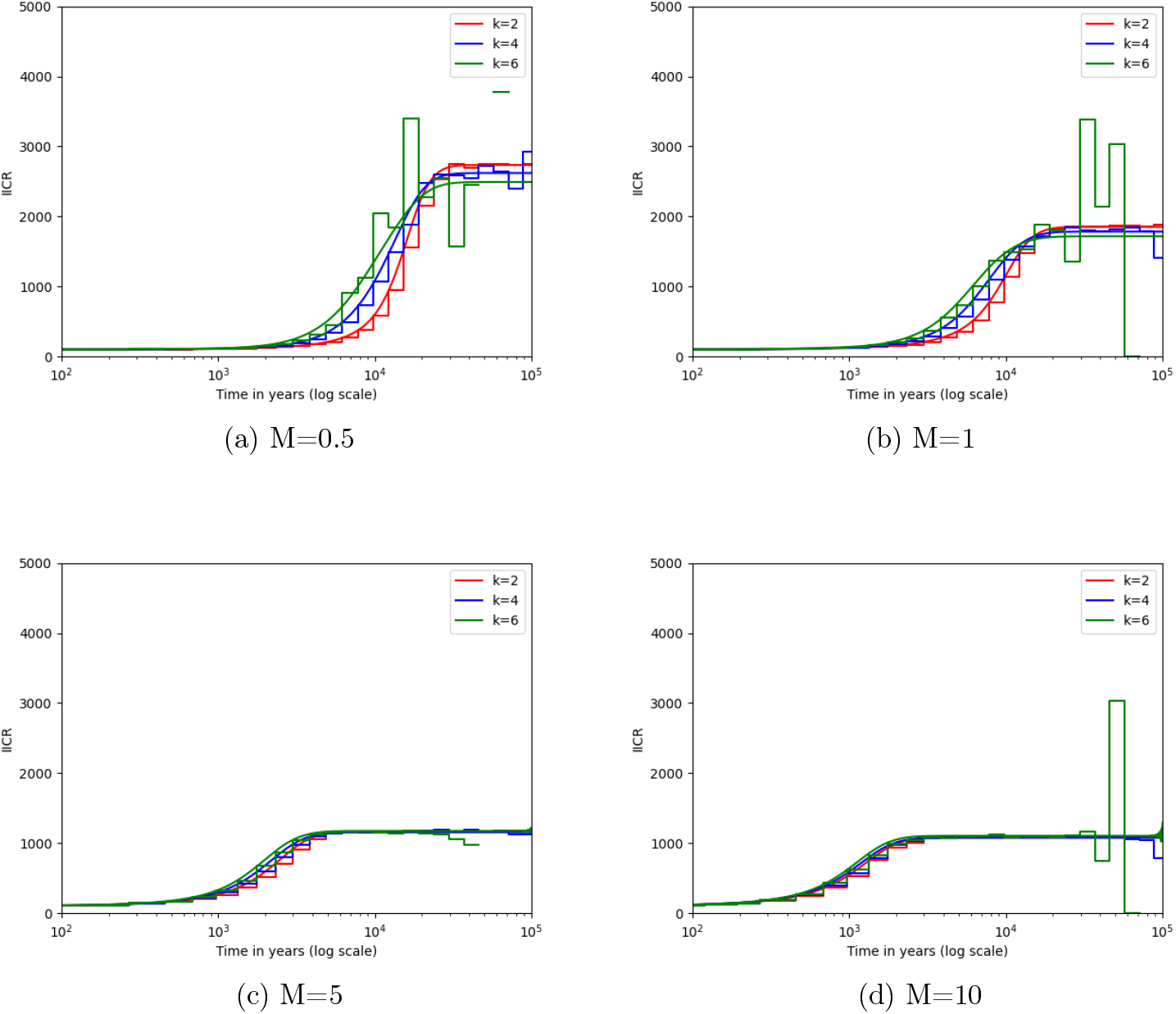
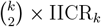 obtained from `IICREstimator` (stepwise curve) and by exact computation (solid curve) for different values of *M*. An *n*-island model were simulated with 10 islands and different migration rates (4*N*_0_*m* = 0.5, 1, 5, 10). All the individuals are sampled in the same deme. 10^6^ *T*_*k*_ were sampled for `IICREstimator`. The curves were then rescaled assuming that *N*_0_ = 100, *g* = 25 and *µ* = 1.25 *×* 10^−8^. The sampling scheme across the ten demes was `[k 0 0 0 0 0 0 0 0 0]` with *k* = 2, 4, 6.

Other sampling schemes were tested where only one diploid individual was sampled per island (Figure S2).

The commands that were used for the different values of *k* were as follow:

~~~
ms 2 1000000 -I 10 2 0 0 0 0 0 0 0 0 0 M -T -L
ms 4 1000000 -I 10 2 2 0 0 0 0 0 0 0 0 M -T -L
ms 6 1000000 -I 10 2 2 2 0 0 0 0 0 0 0 M -T -L
~~~

**Figure S2:**
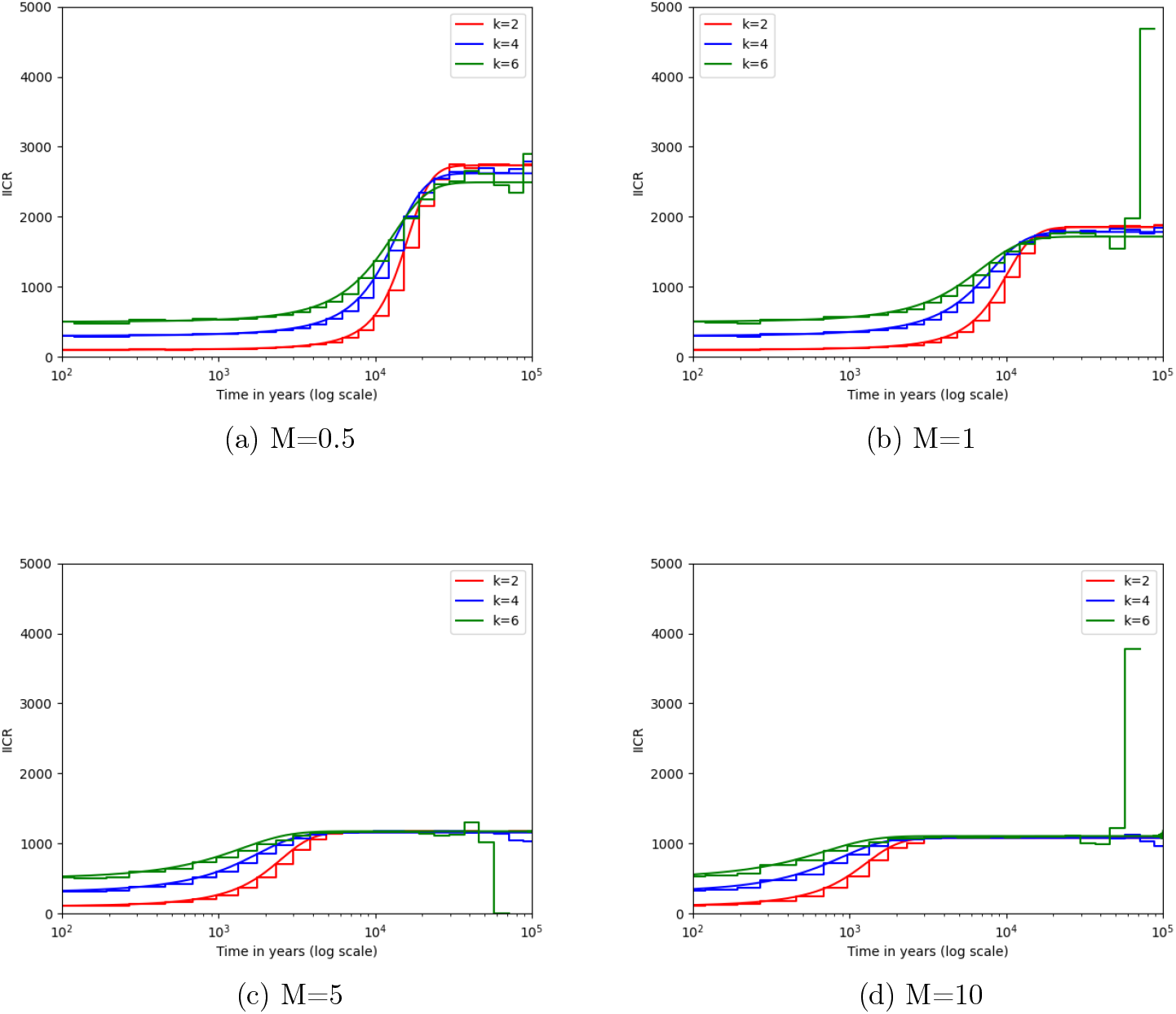
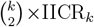 obtained from `IICREstimator` (stepwise curve) and exactly computed (solid curve) for different values of *M*. Each diploid individual was sampled in a different deme. The sampling scheme was `[10 2 0 0 0 0 0 0 0 0 0]` when *k* = 2, `[10 2 2 0 0 0 0 0 0 0 0]` when *k* = 4 and `[10 2 2 2 0 0 0 0 0 0 0]` when *k* = 6.

In addition to the n-island model, a stepping stone model was studied. With this model, the position of the island in which the samples were taken has an effect on the shape of the IICR curves (Figure S4). A simple linear stepping stone with five islands was simulated with the msprime command :

~~~
mspms 4 1 -I 5 4 0 0 0 0 0 -m 1 2 0.5 -m 2 3 0.5 -m 3 4 0.5 -m 4 5 0.5
-m 5 4 0.5 -m 4 3 0.5 -m 3 2 0.5 -m 2 1 0.5 -T
~~~

The diagram (Figure S3) shows the different sampling schemes tested for k=4.

**Figure S3:**
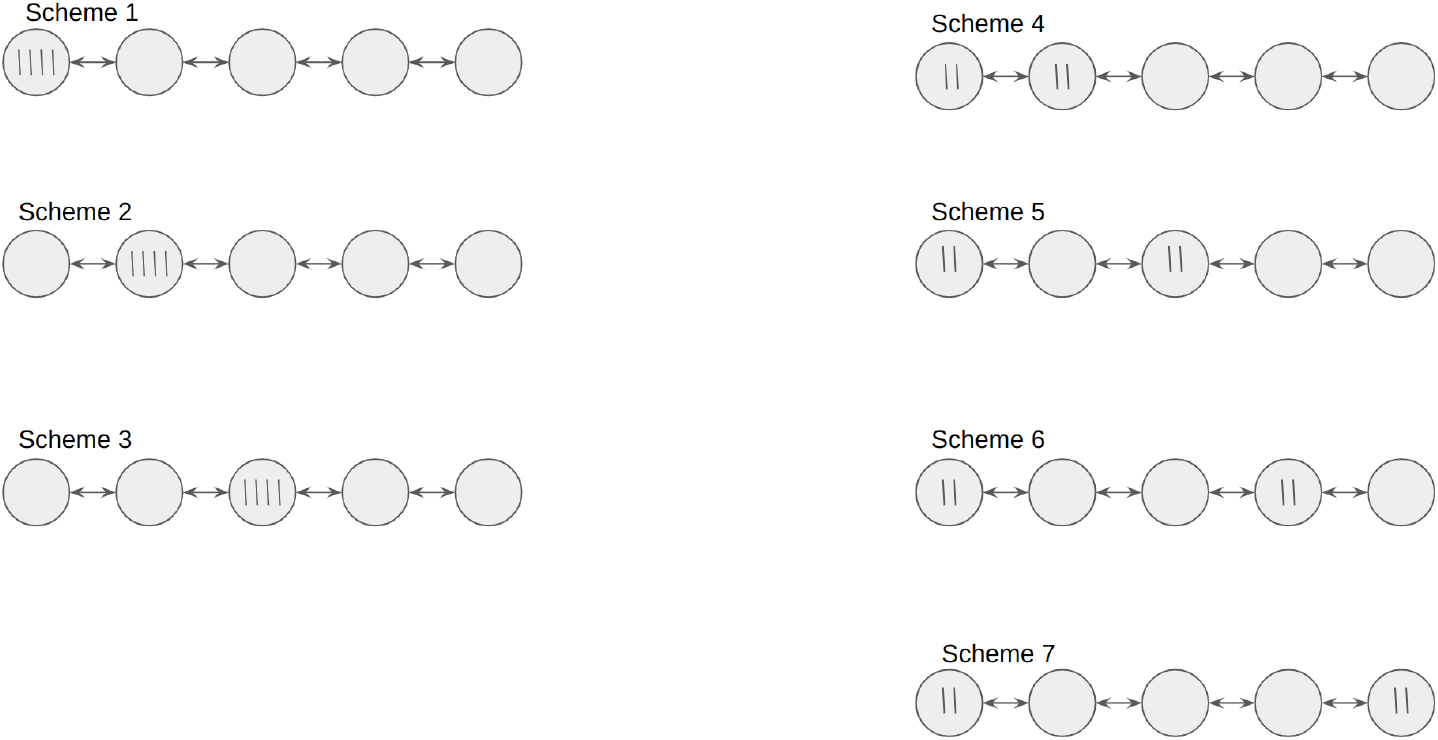
List of the possible sampling schemes for *k* = 4

**Figure S4:**
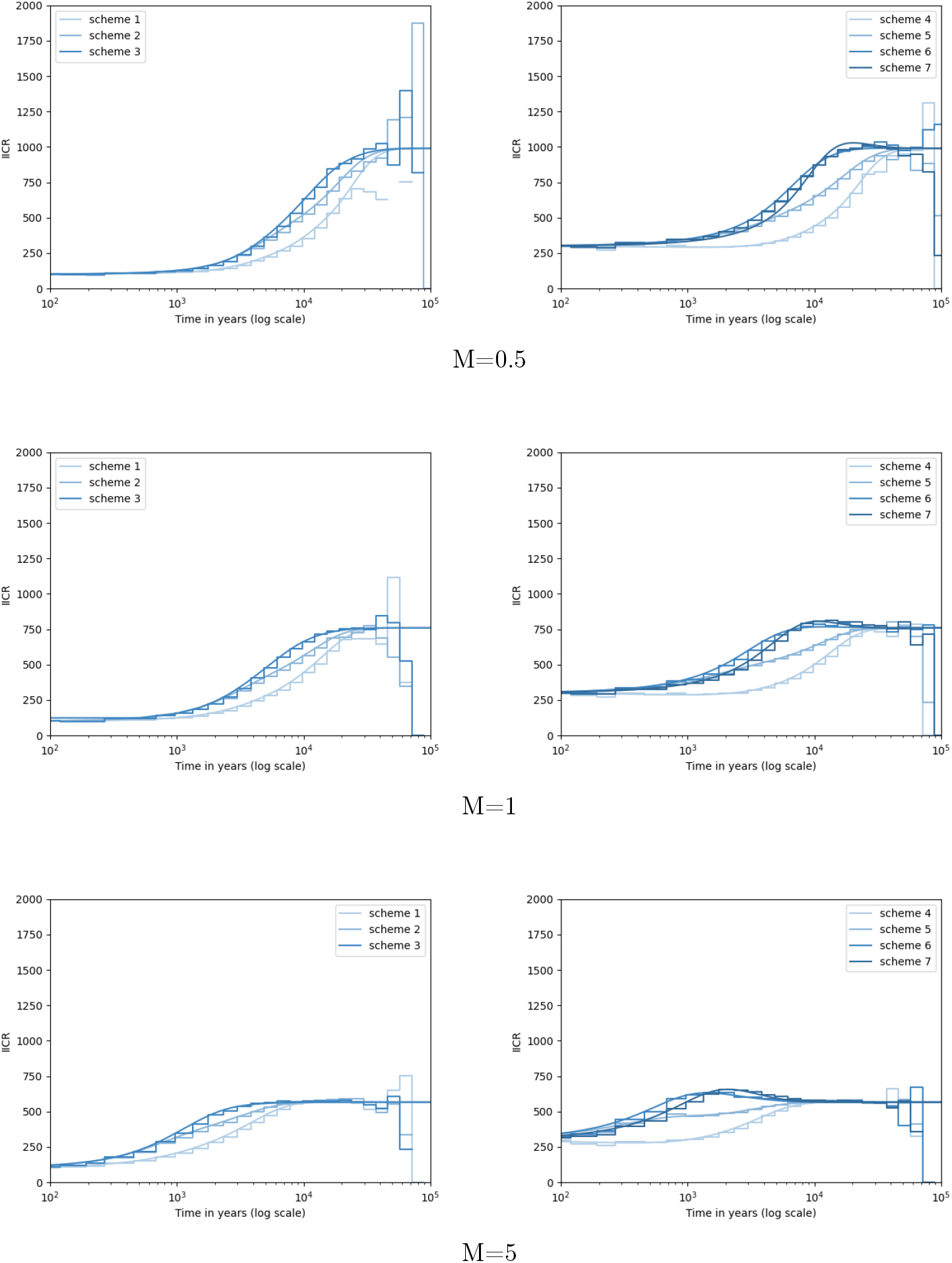
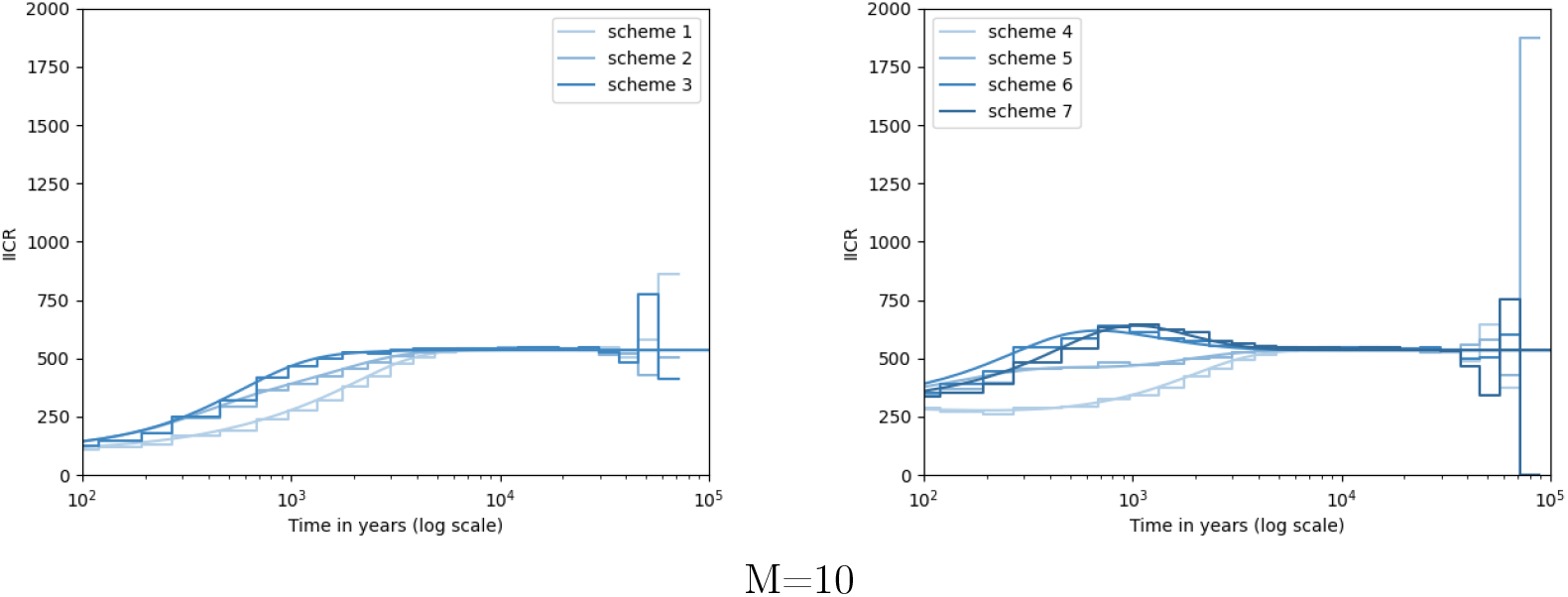
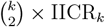 obtained from `IICREstimator` (stepwise curve) and computed with the Q matrix approach (solid curve) for different values of *M* and for the sampling schemes described in Figure S3.

### III. 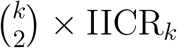 and MSMC

The MSMC method takes several text files as input, each one corresponding to a different chromosome. Here, the genomes were simulated using `msprime` [43]. To obtain a whole genome of 3 *×* 10^9^ BP we simulated ten chromosomes of length 3 *×* 10^8^ BP each. The mutation rate was set to 1.25 *×* 10^−8^ per BP per generation and the recombination rate to 10^−8^ per BP per generation. The maximum number of recombinational segments was set to 8 using the `-p` flag.

Once all the chromosomes had been simulated, the demographic history was computed with MSMC using the following command :

~~~
msmc --fixedRecombination -s -o msmc_results chrom*.txt
~~~

The option `-s` was used to remove unphased sites.

For the sake of consistency, this section has the same structure of the previous one. The first model to be tested is then the *n*-island model where all the individuals were sampled from the same deme (Figure S5). Such scenario can be simulated from the following command:

~~~
mspms k 1 -t 1500 -r 1200 300000000 -I 10 k 0 0 0 0 0 0 0 0 0 M -p 8 -T
~~~

where *k* was set to 2, 4, 6 and *M* to 0.5, 1, 5, 10. The value of `-t` and `-r` were respectively obtained from the equation 4 *× N*_0_ *× µ × l* and 4 *× N* 0 *× ρ × l* where *l* is the length of the chromosome.

**Figure S5:**
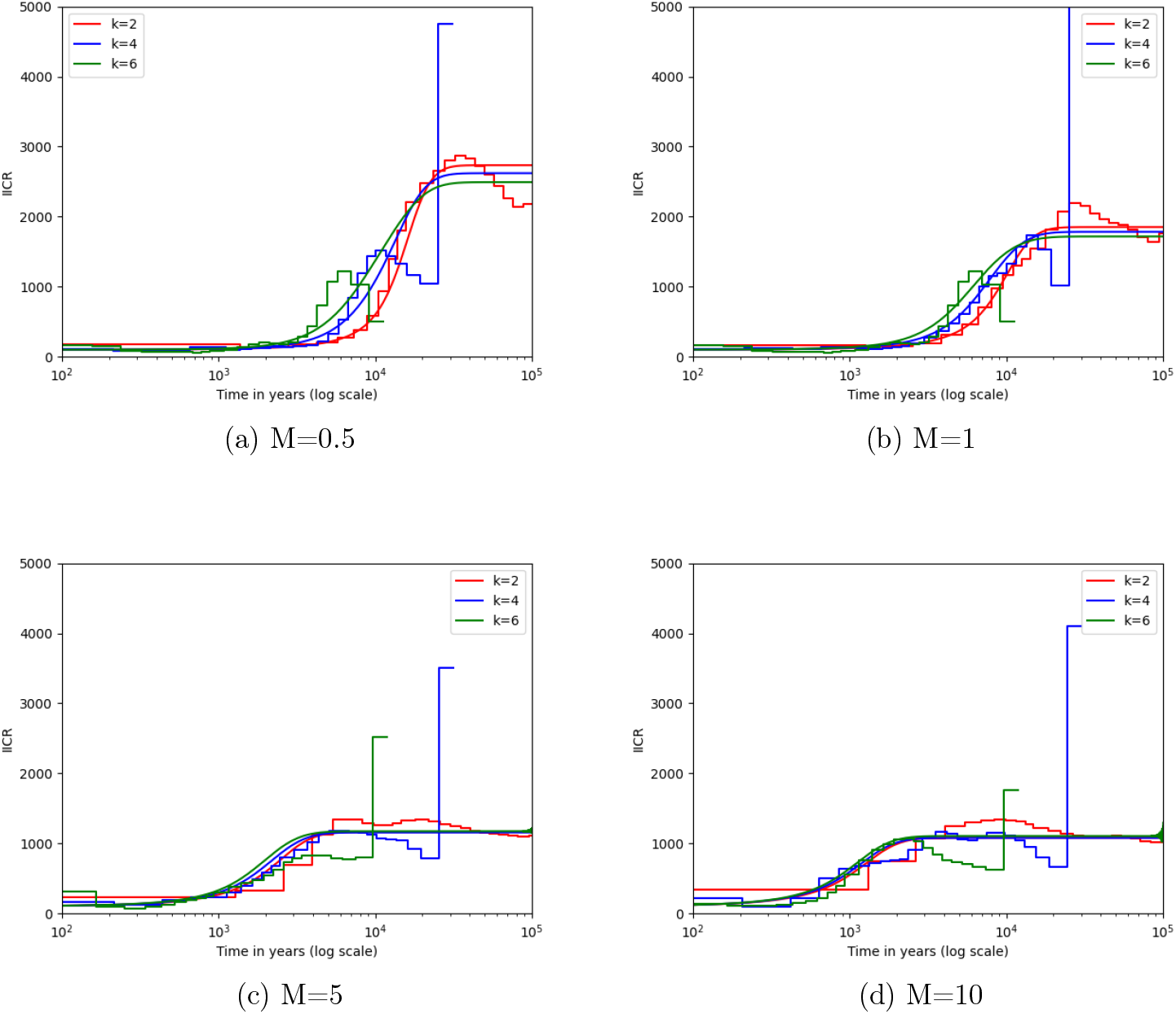
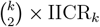 obtained from MSMC (stepwise curve) and exactly computed (solid curve) for different values of *M*. An *n*-island model was simulated with 10 islands and different migration rates (4*N*_0_*m* = 0.5, 1, 5, 10). All the individuals are sampled in the same deme. Genomes of length 3*×*10^9^, recombination rate of 10^−8^ and mutation rate of 1.25*×*10^−8^ were simulated with `msprime`. The size of the deme is 100 and the generation time is 25.

A second sampling scheme was tested where one diploid individual was sampled per deme. In that case, the command was:

~~~
mspms 2 1 -t 1500 -r 1200 300000000 -I 10 2 0 0 0 0 0 0 0 0 0 M -p 8 -T
mspms 4 1 -t 1500 -r 1200 300000000 -I 10 2 2 0 0 0 0 0 0 0 0 M -p 8 -T
mspms 6 1 -t 1500 -r 1200 300000000 -I 10 2 2 2 0 0 0 0 0 0 0 M -p 8 -T
~~~

**Figure S6:**
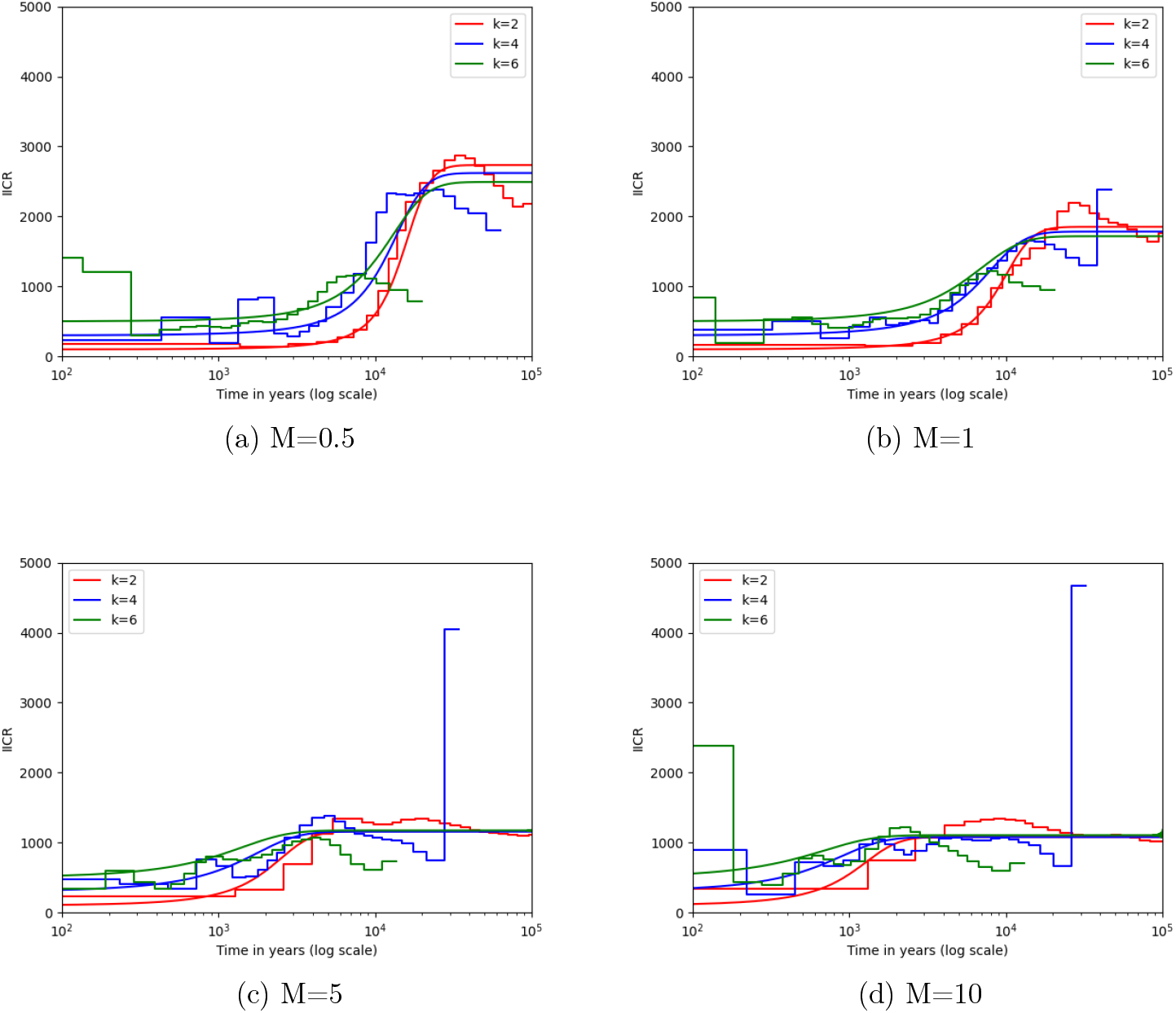
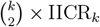 obtained from MSMC (stepwise curve) and computed exactly (solid curve) for different values of *M*. Each individual was sampled in a different deme.

The `msprime` command that was used to generate the genome was :

~~~
mspms 4 1 -t 1500 -r 1200 300000000 -I 5 4 0 0 0 0 0 -m 1 2 0.5 -m 2 3 0.5
-m 3 4 0.5 -m 4 5 0.5 -m 5 4 0.5 -m 4 3 0.5 -m 3 2 0.5 -m 2 1 0.5 -p 8 -T.
~~~

The different sampling schemes used are given in figure S3.

**Figure S7:**
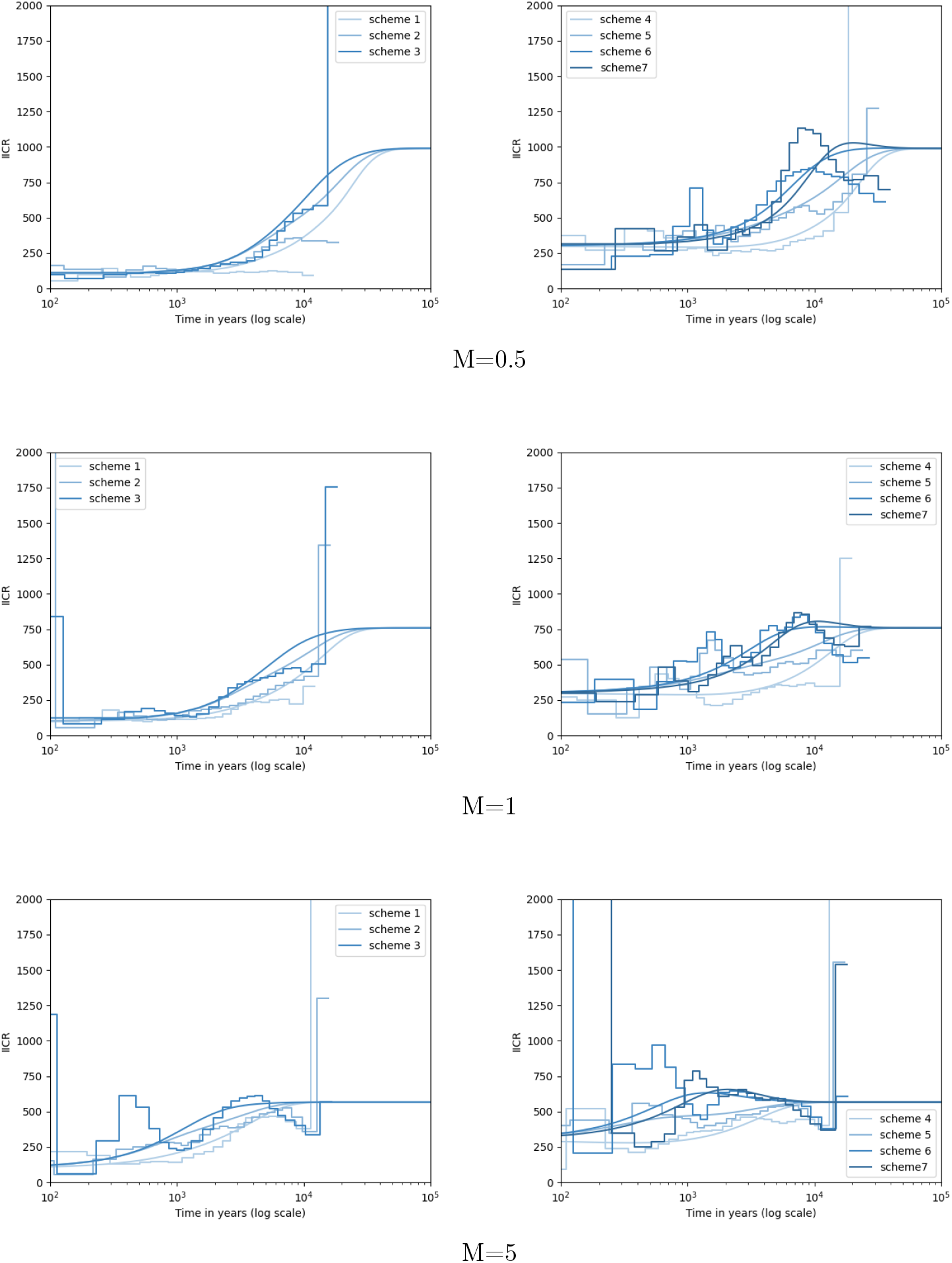
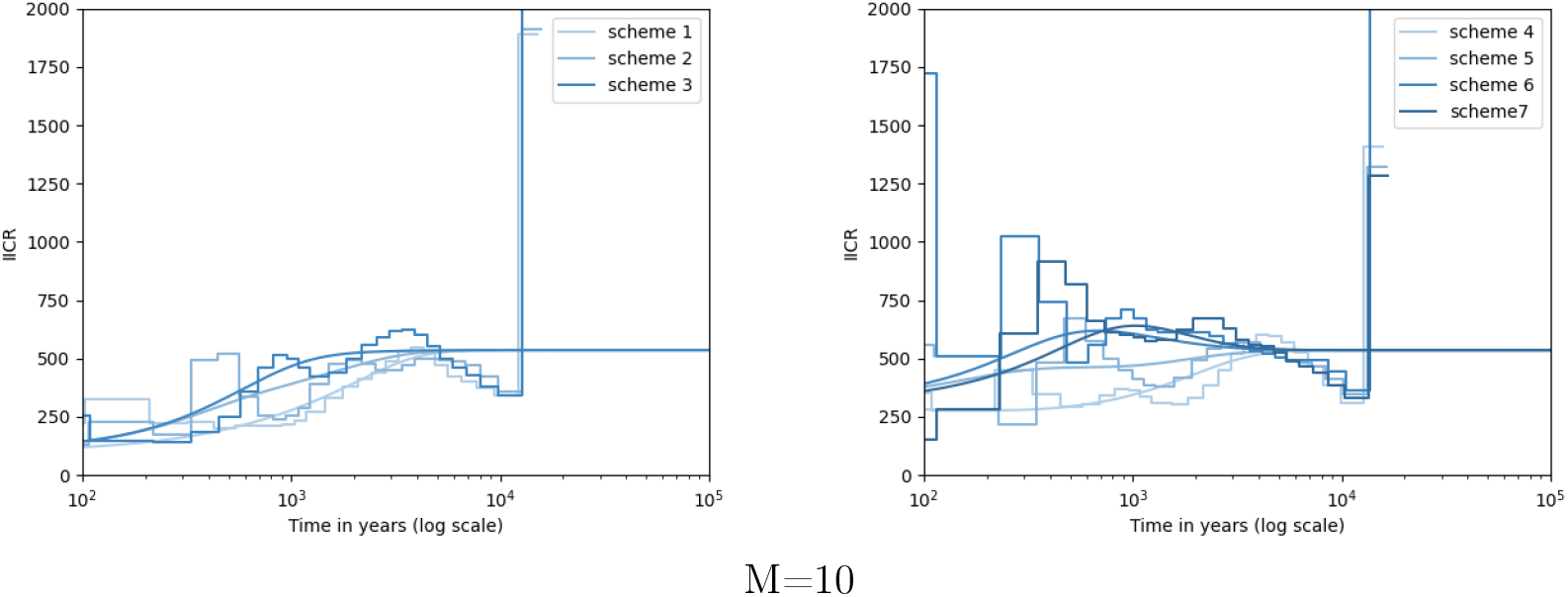
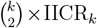 obtained from `IICREstimator` (stepwise curve) and computed exactly (solid curve) for different values of *M* and sampling schemes.

**Figure S8:**
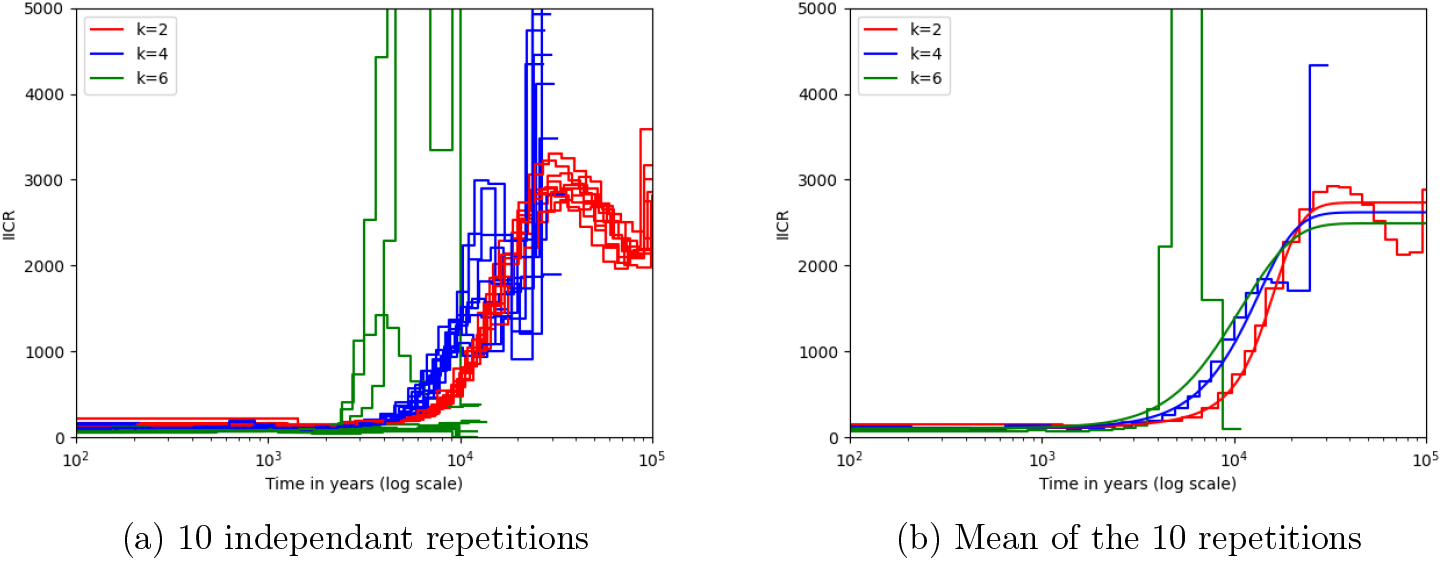
10 independent repetitions of MSMC and the mean curve

Finally, the MSMC method was tested on several independent runs of the same command (10 runs are represented on the figure S8). Only the *n*-island model was tested but different values of *k* were compared. The average of the ten curves were then compared to the theoretical curve (solid line).

### IV. 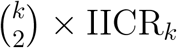 with MSMC2

All the genomes were simulated using `msprime`. The MSMC and MSMC2 methods were applied to the same genome files using the same commands as above. As the PSMC requires only one file and does not need phased genomes, the command used to simulate the genome was :

~~~
mspms 2 10 -t 1500 -r 1200 300000000 -I 10 2 0 0 0 0 0 0 0 0 0 10 -p 8 -T
~~~

**Figure S9:**
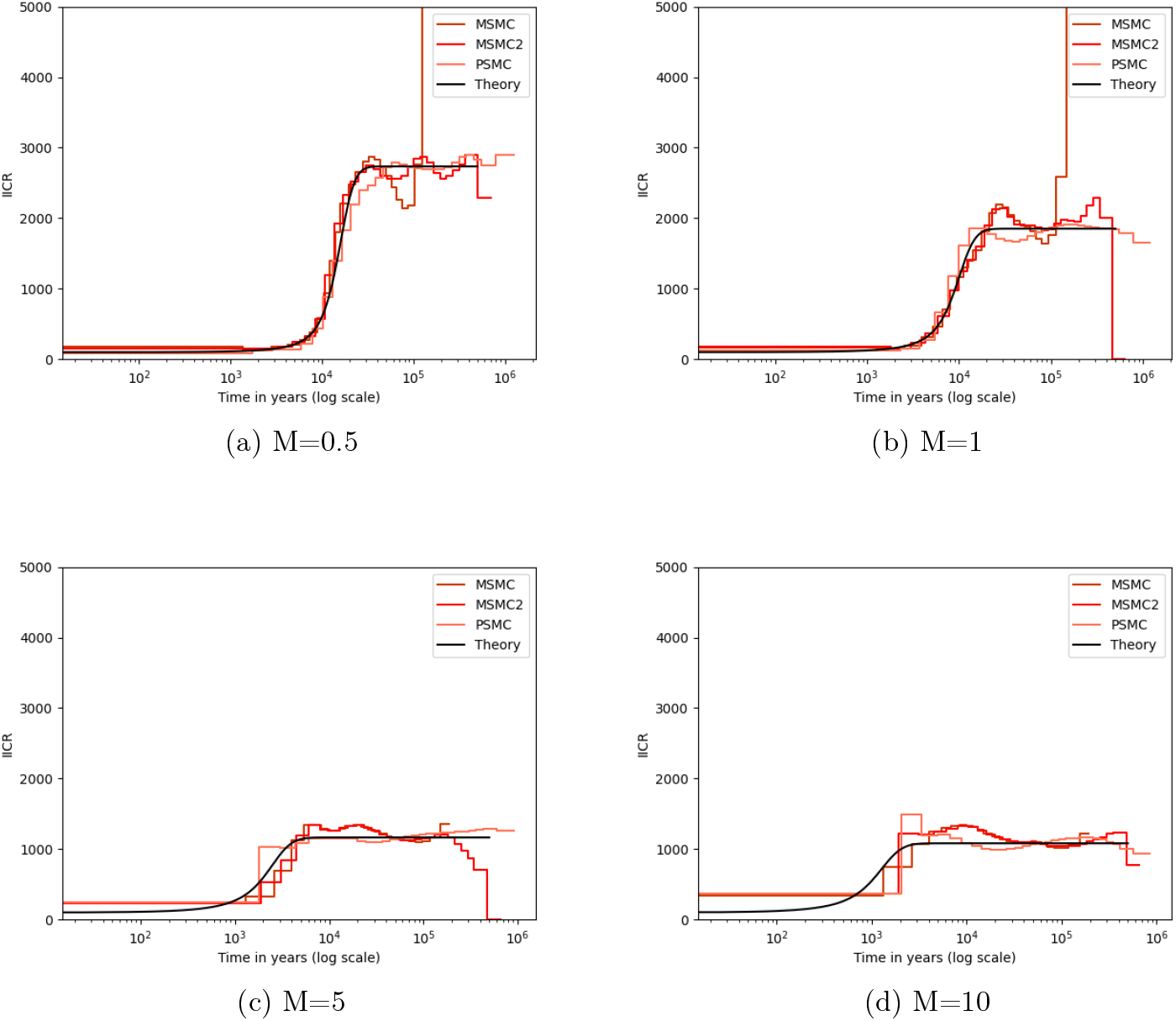
Comparing the PSMC, MSMC and MSMC2 and the theoretical 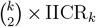. All the individuals were sampled in the same deme taken from a stationary n-island model as defined in the msprime command above

**Figure S10:**
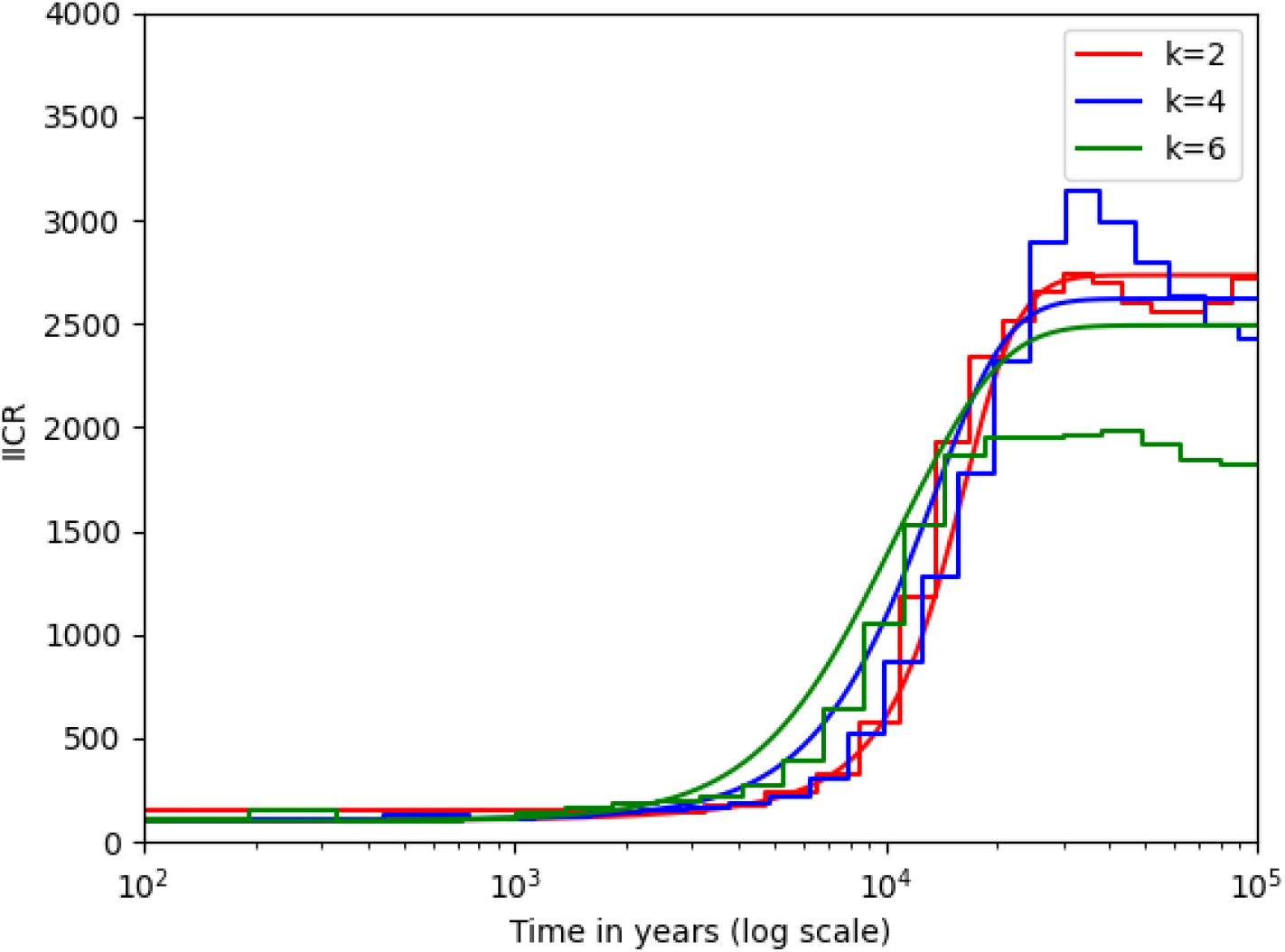
MSMC2 when increasing the sampling size

**Figure S11:**
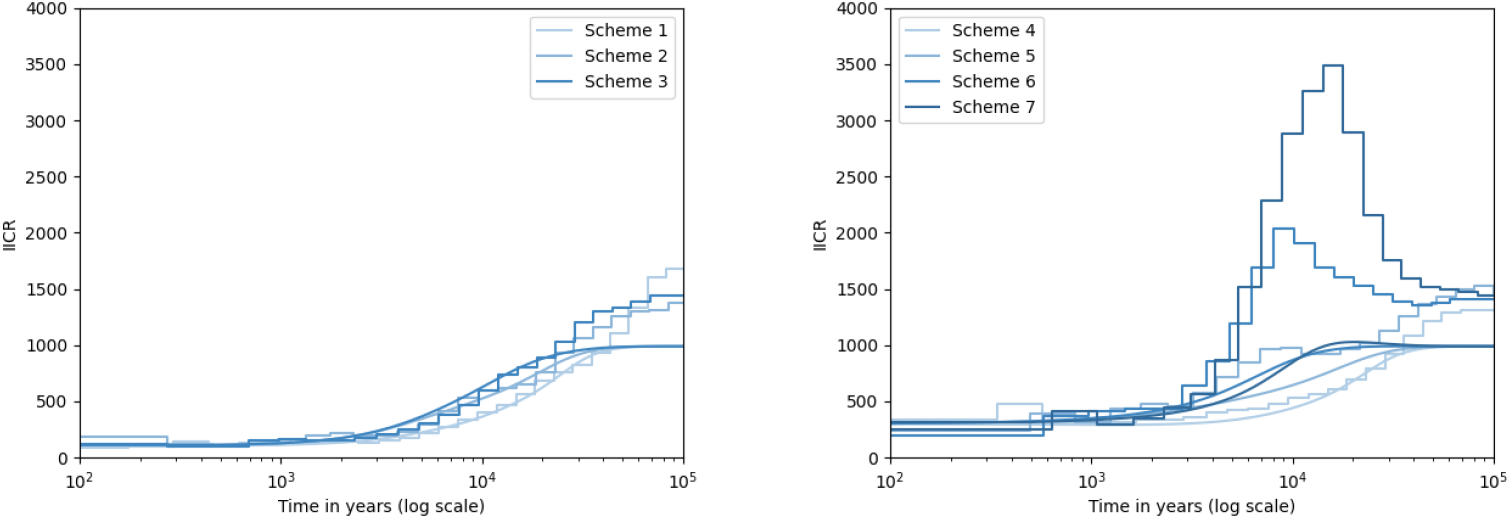
Comparing MSMC2 and the theoretical 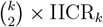 in a Stepping stone model where *M* = 0.5.

### I. IICREstimator parameter file

Example of the parameter file used with `IICREstimator`.

~~~
{
 “path2ms” : “./”,
 “scenarios” : [
 {“ms_command”:”ms 2 1000000 -I 10 2 0 0 0 0 0 0 0 0 0 0.5 -T -L”,
   “label”:”ms”,
   “color”: “r”,
   “linestyle”: “:”,
   “linewidth”: 1,
   “alpha”: 1
   }
 ],
 “theoretical_IICR_general”: [
    ],
 “computation_parameters” : {
    “x_vector_type”: “log”,
    “start”: 0,
    “end”: 100,
    “number_of_values”: 64,
    “pattern”: “4*1+25*2+4+6”
    },
 “custom_x_vector”: {
 “set_custom_xvector”: 0,
 “x_vector”: 0
 },
 “scale_params”:{
 “N0”: 100,
 “generation_time”: 25
 },
 “plot_params”:{
   “plot_theor_IICR”: 0,
   “plot_real_ms_history”: 0,
   “plot_limits”: [1e3, 1e8, 0, 20000],
   “plot_title”: “N-island model”,
   “plot_xlabel”: “time “,
   “plot_ylabel”: “IICR”,
   “show_plot”: 1,
   “save_figure”: 0
 },
 “number_of_repetitions”: {
 “n_rep”: 1
 },
 “vertical_lines”:[],
 “plot_densities”: {
 “densities_to_plot”: [],
 “x_lim”: [0, 600],
 “y_lim”: [0, 600]
 },
 “save_IICR_as_file”:1
}
~~~

### II. Python code for plots

~~~
#extract time and IICR values from tk sampling
def fonctionT2(file):
    x=[]
    y=[]
    a=open(file,’r’) lines=a.read() a.close()
    x_lines=(lines.split(‘\n’)[0])
    x_lines=(x_lines.split(‘ ‘))
    y_lines=(lines.split(‘\n’)[1])
    y_lines=(y_lines.split(‘ ‘))
    for i in range(len(x_lines)):
        x.append(float(x_lines[i]))
        y.append(float(y_lines[i]))
    return(x,y)
#extract time and IICR values from Q matrix
def fonctionTheo(file, N, g): #N is the size of the deme and g is the generation time
    x=[]
    y=[]
    a=open(file,’r’)
    lines=a.read()
    a.close()
    lines=lines.split(‘\n’)
    x_lines=lines[-1].split(‘ ‘)
    y_lines=lines[0:-1]
    for i in range(len(y_lines)):
        x.append(float(x_lines[i]))
        y.append(float(y_lines[i]))
    times = [g * 2 * N * i for i in x]
    sizes = [N * i for i in y]
    return(times,sizes)
~~~

## References

[1] Adams, D. (1979). The Hitchhiker’s Guide to the Galaxy. Pan Books.

[2] Alcala, N. and Vuilleumier, S. (2014). Turnover and accumulation of genetic diversity across large time-scale cycles of isolation and connection of populations. Proceedings of the Royal Society B: Biological Sciences, 281(1794):20141369.

[3] Arenas, M., Ray, N., Currat, M., and Excoffier, L. (2012). Consequences of range contractions and range shifts on molecular diversity. Molecular biology and evolution, 29(1):207–218.

[4] Arredondo, A., Mourato, B., Nguyen, K., Boitard, S., Rodríguez, W., Mazet, O., and Chikhi, L. (2021). Inferring number of populations and changes in connectivity under the n-island model. Heredity, 126(6):896–912.

[5] Balboa, R. F., Bertola, L. D., Brüniche-Olsen, A., Rasmussen, M. S., Liu, X., Besnard, G., Salmona, J., Santander, C. G., He, S., Zinner, D., et al. (2024). African bushpigs exhibit porous species boundaries and appeared in madagascar concurrently with human arrival. Nature Communications, 15(1):172.

[6] Beaumont, M. (2004). Recent developments in genetic data analysis: what can they tell us about human demographic history? Heredity, 92(5):365–379.

[7] Beaumont, M. A. (1999). Detecting population expansion and decline using microsatellites. Genetics, 153(4):2013–2029.

[8] Beaumont, M. A. (2010). Approximate Bayesian computation in evolution and ecology. Annual Review of Ecology, Evolution, and Systematics, 41:379–406.

[9] Beaumont, M. A., Zhang, W., and Balding, D. J. (2002). Approximate Bayesian computation in population genetics. Genetics, 162(4):2025–2035.

[10] Beichman, A. C., Phung, T. N., and Lohmueller, K. E. (2017). Comparison of single genome and allele frequency data reveals discordant demographic histories. G3: Genes, Genomes, Genetics, 7(11):3605–3620.

[11] Bergman, J., Pedersen, R. Ø., Lundgren, E. J., Lemoine, R. T., Monsarrat, S., Pearce, E. A., Schierup, M. H., and Svenning, J.-C. (2023). Worldwide late pleistocene and early holocene population declines in extant megafauna are associated with homo sapiens expansion rather than climate change. Nature Communications, 14(1):7679.

[12] Boitard, S., Arredondo, A., Chikhi, L., and Mazet, O. (2022). Heterogeneity in effective size across the genome: effects on the inverse instantaneous coalescence rate (iicr) and implications for demographic inference under linked selection. Genetics, 220(3):iyac008.

[13] Boitard, S., Rodríguez, W., Jay, F., Mona, S., and Austerlitz, F. (2016). Inferring population size history from large samples of genome-wide molecular data - an approximate Bayesian computation approach. PLoS Genetics, 12(3):e1005877.

[14] Brand, C. M., White, F. J., Rogers, A. R., and Webster, T. H. (2022). Estimating bonobo (Pan paniscus) and chimpanzee (Pan troglodytes) evolutionary history from nucleotide site patterns. Proceedings of the National Academy of Sciences of the United States of America, 119(17):1–12.

[15] Bunnefeld, L., Frantz, L. A., and Lohse, K. (2015). Inferring bottlenecks from genomewide samples of short sequence blocks. Genetics, 201(3):1157–1169.

[16] Chikhi, L., Rodríguez, W., Grusea, S., Santos, P., Boitard, S., and Mazet, O. (2018). The IICR (inverse instantaneous coalescence rate) as a summary of genomic diversity: insights into demographic inference and model choice. Heredity, 120:13–24.

[17] Chikhi, L., Sousa, V. C., Luisi, P., Goossens, B., and Beaumont, M. A. (2010). The confounding effects of population structure, genetic diversity and the sampling scheme on the detection and quantification of population size changes. Genetics, 186(3):983–995.

[18] Cranmer, K., Brehmer, J., and Louppe, G. (2020). The frontier of simulation-based inference. Proceedings of the National Academy of Sciences, 117(48):30055–30062.

[19] Eriksson, A. and Manica, A. (2012). Effect of ancient population structure on the degree of polymorphism shared between modern human populations and ancient hominins. Proceedings of the National Academy of Sciences, 109(35):13956–13960.

[20] Eriksson, A. and Manica, A. (2014). The doubly conditioned frequency spectrum does not distinguish between ancient population structure and hybridization. Molecular Biology and Evolution, 31(6):1618–1621.

[21] Excoffier, L. (2004). Patterns of DNA sequence diversity and genetic structure after a range expansion: lessons from the infinite-island model. Molecular ecology, 13(4):853–864.

[22] Excoffier, L., Dupanloup, I., Huerta-Sánchez, E., Sousa, V. C., and Foll, M. (2013). Robust demographic inference from genomic and snp data. PLoS Genetics, 9(10):e1003905.

[23] Fitak, R. R., Mohandesan, E., Corander, J., and Burger, P. A. (2016). The de novo genome assembly and annotation of a female domestic dromedary of north african origin. Molecular ecology resources, 16(1):314–324.

[24] Godoy, J. A., Negro, J. J., Hiraldo, F., and Donázar, J. A. (2004). Phylogeography, genetic structure and diversity in the endangered bearded vulture (gypaetus barbatus, l.) as revealed by mitochondrial dna. Molecular Ecology, 13(2):371–390.

[25] Goldstein, D. B. and Chikhi, L. (2002). Human migrations and population structure: what we know and why it matters. Annual Review of Genomics and Human Genetics, 3(1):129–152.

[26] Goossens, B., Chikhi, L., Ancrenaz, M., Lackman-Ancrenaz, I., Andau, P., Bruford, M. W., et al. (2006). Genetic signature of anthropogenic population collapse in orangutans. PLoS Biology, 4(2):285.

[27] Griffiths, R. and Tavaré, S. (1994). Simulating probability distributions in the coalescent. Theoretical Population Biology, 46(2):131–159.

[28] Groenen, M., Archibald, A., Uenishi, H., Tuggle, C., Takeuchi, Y., Rothschild, M., Rogel-Gaillard, C., Park, C., Milan, D., Megens, H., Li, S., Larkin, D., Kim, H., Frantz, L., Caccamo, M., Ahn, H., Aken, B., Anselmo, A., Anthon, C., Auvil, L., Badaoui, B., Beattie, C., Bendixen, C., Berman, D., Blecha, F., Blomberg, J., Bolund, L., Bosse, M., Botti, S., and Bujie, Z. (2012). Analyses of pig genomes provide insight into porcine demography and evolution. Nature, 491:393–398.

[29] Grusea, S., Rodríguez, W., Pinchon, D., Chikhi, L., Boitard, S., and Mazet, O. (2018). Coalescence times for three genes provide sufficient information to distinguish population structure from population size changes. Journal of Mathematical Biology, 78(1-2):189–224.

[30] Guevara, E. E., Webster, T. H., Lawler, R. R., Bradley, B. J., Greene, L. K., Ranaivonasy, J., Ratsirarson, J., Harris, R. A., Liu, Y., Murali, S., Raveendran, M., Hughes, D. S. T., Muzny, D. M., Yoder, A. D., Worley, K. C., and Rogers, J. (2021). Comparative genomic analysis of sifakas (Propithecus) reveals selection for folivory and high heterozygosity despite endangered status. Science advances, 7.

[31] Gutenkunst, R. N., Hernandez, R. D., Williamson, S. H., and Bustamante, C. D. (2009). Inferring the joint demographic history of multiple populations from multidimensional SNP frequency data. PLoS Genetics, 5(10):e1000695.

[32] Heller, R., Chikhi, L., and Siegismund, H. R. (2013). The confounding effect of population structure on Bayesian skyline plot inferences of demographic history. PLoS One, 8(5):e62992.

[33] Herbots, H. M. J. D. (1994). Stochastic models in population genetics: genealogy and genetic differentiation in structured populations. PhD thesis.

[34] Hewitt, G. (2000). The genetic legacy of the Quaternary ice ages. Nature, 405(6789):907–913.

[35] Hewitt, G. M. (1996). Some genetic consequences of ice ages, and their role in divergence and speciation. Biological Journal of the Linnean Society, 58(3):247–276.

[36] Hey, J. and Machado, C. A. (2003). The study of structured populations–new hope for a difficult and divided science. Nature Reviews. Genetics, 4(7):535.

[37] Hudson, R. R. (1990). Gene genealogies and the coalescent process. Oxford surveys in evolutionary biology, 7(1):44.

[38] Hudson, R. R. (2002). Generating samples under a Wright–Fisher neutral model of genetic variation. Bioinformatics, 18(2):337–338.

[39] Johri, P., Aquadro, C. F., Beaumont, M., Charlesworth, B., Excoffier, L., Eyre-Walker, A., Keightley, P. D., Lynch, M., McVean, G., Payseur, B. A., et al. (2022). Recommendations for improving statistical inference in population genomics. PLoS biology, 20(5):e3001669.

[40] Johri, P., Charlesworth, B., and Jensen, J. D. (2020). Toward an evolutionarily appropriate null model: jointly inferring demography and purifying selection. Genetics, 215(1):173–192.

[41] Kamm, J., Terhorst, J., Durbin, R., and Song, Y. S. (2019). Efficiently inferring the demographic history of many populations with allele count data. Journal of the American Statistical Association, pages 1–16.

[42] Kelleher, J., Etheridge, A. M., and McVean, G. (2016). Efficient coalescent simulation and genealogical analysis for large sample sizes. PLoS computational biology, 12(5):e1004842.

[43] Kelleher, J. and Lohse, K. (2020). Coalescent Simulation with msprime. Methods in Molecular Biology. Springer US, New York, NY.

[44] Li, H. and Durbin, R. (2011). Inference of human population history from individual whole-genome sequences. Nature, 475(7357):493–496.

[45] Liu, S., Simons, J., Yi, M., and Beaumont, M. (2023). Variational gradient descent using local linear models. arXiv preprint arXiv:2305.15577.

[46] Liu, X. and Fu, Y.-X. (2015). Exploring population size changes using SNP frequency spectra. Nature genetics.

[47] Lueckmann, J.-M., Boelts, J., Greenberg, D., Goncalves, P., and Macke, J. (2021). Benchmarking simulation-based inference. In International conference on artificial intelligence and statistics, pages 343–351. PMLR.

[48] Malaspinas, A.-S., Westaway, M. C., Muller, C., Sousa, V. C., Lao, O., Alves, I., Bergström, A., Athanasiadis, G., Cheng, J. Y., Crawford, J. E., et al. (2016). A genomic history of aboriginal australia. Nature, 538(7624):207–214.

[49] Mazet, O., Rodríguez, W., and Chikhi, L. (2015). Demographic inference using genetic data from a single individual: Separating population size variation from population structure. Theoretical Population Biology, 104:46–58.

[50] Mazet, O., Rodríguez, W., Grusea, S., Boitard, S., and Chikhi, L. (2016). On the importance of being structured: instantaneous coalescence rates and human evolution— lessons for ancestral population size inference? Heredity, 116(4):362.

[51] Miller, W., Drautz, D. I., Janecka, J. E., Lesk, A. M., Ratan, A., Tomsho, L. P., Packard, M., Zhang, Y., McClellan, L. R., Qi, J., et al. (2009). The mitochondrial genome sequence of the tasmanian tiger (thylacinus cynocephalus). Genome research, 19(2):213–220.

[52] Nielsen, R. and Beaumont, M. A. (2009). Statistical inferences in phylogeography. Molecular Ecology, 18(6):1034–1047.

[53] [Paris] Paris, C. Modélisation de la généalogie d’une population structurée. Master’s thesis, Institut National de Sciences Appliquées de Toulouse.

[54] Parreira, B. R. and Chikhi, L. (2015). On some genetic consequences of social structure, mating systems, dispersal, and sampling. Proceedings of the National Academy of Sciences, 112(26):E3318–E3326.

[55] Paz-Vinas, I., Quéméré, E., Chikhi, L., Loot, G., and Blanchet, S. (2013). The demographic history of populations experiencing asymmetric gene flow: combining simulated and empirical data. Molecular Ecology, 22(12):3279–3291.

[56] Peter, B. M., Wegmann, D., and Excoffier, L. (2010). Distinguishing between population bottleneck and population subdivision by a Bayesian model choice procedure. Molecular Ecology, 19(21):4648–4660.

[57] Poelstra, J. W., Salmona, J., Tiley, G. P., Schü ßler, D., Blanco, M. B., Andriambeloson, J. B., Bouchez, O., Campbell, C. R., Etter, P. D., Hohenlohe, P. A., Hunnicutt, K. E., Iribar, A., Johnson, E. A., Kappeler, P. M., Larsen, P. A., Manzi, S., Ralison, J. M., Randrianambinina, B., Rasoloarison, R. M., Rasolofoson, D. W., Stahlke, A. R., Weisrock, D. W., Williams, R. C., Chikhi, L., Louis, E. E., Radespiel, U., and Yoder, A. D. (2021). Cryptic patterns of speciation in cryptic primates: Microendemic mouse lemurs and the multispecies coalescent. Systematic biology, 70:203–218.

[58] Quéméré, E., Amelot, X., Pierson, J., Crouau-Roy, B., and Chikhi, L. (2012). Genetic data suggest a natural prehuman origin of open habitats in northern Madagascar and question the deforestation narrative in this region. Proceedings of the National Academy of Sciences, 109(32):13028–13033.

[59] Rasteiro, R., Bouttier, P.-A., Sousa, V. C., and Chikhi, L. (2012). Investigating sexbiased migration during the neolithic transition in europe, using an explicit spatial simulation framework. Proceedings. Biological sciences, 279:2409–2416.

[60] Raxworthy, C. J. and Smith, B. T. (2021). Mining museums for historical dna: advances and challenges in museomics. Trends in Ecology & Evolution, 36(11):1049–1060.

[61] Rodríguez, W., Mazet, O., Grusea, S., Arredondo, A., Corujo, J. M., Boitard, S., and Chikhi, L. (2018). The IICR and the non-stationary structured coalescent: towards demographic inference with arbitrary changes in population structure. Heredity, 121(6):663–678.

[62] Rogers, A. R. and Harpending, H. (1992). Population growth makes waves in the distribution of pairwise genetic differences. Molecular biology and evolution, 9(3):552–569.

[63] Salmona, J., Heller, R., Quéméré, E., and Chikhi, L. (2017). Climate change and human colonization triggered habitat loss and fragmentation in madagascar. Molecular Ecology, 26(19):5203–5222.

[64] Scerri, E. M. L., Thomas, M. G., Manica, A., Gunz, P., Stock, J. T., Stringer, C., Grove, M., Groucutt, H. S., Timmermann, A., Rightmire, G. P., d’Errico, F., Tryon, C. A., Drake, N. A., Brooks, A. S., Dennell, R. W., Durbin, R., Henn, B. M., Lee-Thorp, J., deMenocal, P., Petraglia, M. D., Thompson, J. C., Scally, A., and Chikhi, L. (2018). Did our species evolve in subdivided populations across Africa, and why does it matter? Trends in Ecology & Evolution.

[65] Schiffels, S. and Durbin, R. (2013). Inferring human population size and separation history from multiple genome sequences. Nature Genetics, 8(46):919–925.

[66] Schiffels, S. and Wang, K. (2020). Msmc and msmc2: the multiple sequentially Markovian coalescent. In Statistical population genomics, pages 147–165. Humana.

[67] Schneider, N., Chikhi, L., Currat, M., and Radespiel, U. (2010). Signals of recent spatial expansions in the grey mouse lemur (microcebus murinus). BMC Evolutionary Biology, 10(1):1–17.

[68] Simon, A. and Coop, G. (2023). The contribution of gene flow, selection, and genetic drift to five thousand years of human allele frequency change. bioRxiv, pages 2023–07.

[69] Sjödin, P., Kaj, I., Krone, S., Lascoux, M., and Nordborg, M. (2005). On the meaning and existence of an effective population size. Genetics, 169(2):1061–1070.

[70] Slatkin, M. and Hudson, R. R. (1991). Pairwise comparisons of mitochondrial DNA sequences in stable and exponentially growing populations. Genetics, 129(2):555–562.

[71] Speidel, L., Forest, M., Shi, S., and Myers, S. R. (2019). A method for genome-wide genealogy estimation for thousands of samples. Nature genetics, 51(9):1321–1329.

[72] Städler, T., Haubold, B., Merino, C., Stephan, W., and Pfaffelhuber, P. (2009). The impact of sampling schemes on the site frequency spectrum in nonequilibrium subdivided populations. Genetics, 182(1):205–216.

[73] Tajima, F. (1989). The effect of change in population size on DNA polymorphism. Genetics, 123(3):597–601.

[74] Teixeira, H., Montade, V., Salmona, J., Metzger, J., Bremond, L., Kasper, T., Daut, G., Rouland, S., Ranarilalatiana, S., Rakotondravony, R., Chikhi, L., Behling, H., and Radespiel, U. (2021a). Past environmental changes affected lemur population dynamics prior to human impact in madagascar. Communications biology, 4:1084.

[75] Teixeira, H., Salmona, J., Arredondo, A., Mourato, B., Manzi, S., Rakotondravony, R., Mazet, O., Chikhi, L., Metzger, J., and Radespiel, U. (2021b). Impact of model assumptions on demographic inferences: the case study of two sympatric mouse lemurs in northwestern madagascar. BMC ecology and evolution, 21:197.

[76] Terhorst, J., Kamm, J. A., and Song, Y. S. (2017). Robust and scalable inference of population history from hundreds of unphased whole genomes. Nature genetics, 49(2):303– 309.

[77] Tournebize, R. and Chikhi, L. (2023). Questioning neanderthal admixture: on models, robustness and consensus in human evolution. bioRxiv, pages 2023–04.

[78] Vishwakarma, R., Sgarlata, G. M., Soriano-Panos, D., Rasteiro, R., Maie, T., Paixao, T., Tournebize, R., and Chikhi, L. (2024). Life history traits influence the dynamics of genetic diversity in a refugium population undergoing expansion and contraction. bioRxiv.

[79] Wakeley, J. (1999). Nonequilibrium migration in human history. Genetics, 153(4):1863– 1871.

[80] Wakeley, J. (2001). The coalescent in an island model of population subdivision with variation among demes. Theoretical population biology, 59(2):133–144.

[81] Wang, K., Mathieson, I., O’Connell, J., and Schiffels, S. (2020). Tracking human population structure through time from whole genome sequences. PLoS Genetics, 16(3):e1008552.

[82] Ward, D., Cannon, P., Beaumont, M., Fasiolo, M., and Schmon, S. (2022). Robust neural posterior estimation and statistical model criticism. Advances in Neural Information Processing Systems, 35:33845–33859.

[83] Williams, R. C., Blanco, M. B., Poelstra, J. W., Hunnicutt, K. E., Comeault, A. A., and Yoder, A. D. (2020). Conservation genomic analysis reveals ancient introgression and declining levels of genetic diversity in madagascar’s hibernating dwarf lemurs. Heredity, 124:236–251.

[84] Willis, K. J., Bennett, K. D., Walker, D., and Hewitt, G. M. (2004). Genetic consequences of climatic oscillations in the Quaternary. Philosophical Transactions of the Royal Society of London. Series B: Biological Sciences, 359(1442):183–195.

[85] Yang, M. A. (2022). A genetic history of migration, diversification, and admixture in asia. Human Population Genetics and Genomics, 2(1).

[86] Zhan, X., Pan, S., Wang, J., Dixon, A., He, J., Muller, M. G., Ni, P., Hu, L., Liu, Y., Hou, H., et al. (2013). Peregrine and saker falcon genome sequences provide insights into evolution of a predatory lifestyle. Nature Genetics, 45(5):563–566.

[87] Zhao, S., Zheng, P., Dong, S., Zhan, X., Wu, Q., Guo, X., Hu, Y., He, W., Zhang, S., Fan, W., et al. (2013). Whole-genome sequencing of giant pandas provides insights into demographic history and local adaptation. Nature Genetics, 45(1):67–71.

